# Strain-level diversity shapes competitive outcomes in *Fusarium*: a multi-omic synthesis of fungal warfare

**DOI:** 10.64898/2026.08.17.744998

**Authors:** M. Navarro, F. Dumetz, E. Groppi, M. Vansteelandt, A. Gadea, M. Haddad, N. Mach, N. Ponts

## Abstract

Fusarium head blight (FHB) is driven by co-occurring *Fusarium* species. Yet the molecular bases of their competitive interactions, particularly at the strain level, remain largely unknown. We performed an integrated multi-omic investigation of four *Fusarium* isolates cultivated in monoculture, self-confrontation (SC) and inter-specific confrontation (C) assays: two *Fusarium graminearum* strains FgrI349 and FgrPH-1, and two *Fusarium avenaceum* strains FaveI494 and FaLH03. Light microscopy and quantitative colorimetry revealed marked phenotypic heterogeneity. the *F. graminearum* strains formed expansive, red-pigmented colonies with rapid radial growth, whereas the *F. avenaceum* isolates grew more slowly and displayed distinct colony morphologies and pigmentation patterns. Untargeted LC-HRMS detected 1,008 metabolites in monocultures and 938 metabolites in confrontation zones. Species-level chemical signatures were confirmed, and strain-specific metabolite sets were identified, with FaLH03 producing more than 60 % of the metabolites being made exclusively by a single strain, highlighting its exceptionally unique metabolic profile.

RNA-seq uncovered extensive transcriptional reprogramming during competition. In self-confrontations, strain-specific differences persisted but no major morphological or metabolic shifts were observed. Inter-specific confrontations elicited partner-dependent responses: FgrI349 up-regulated 1,492 genes against FaveI494 (including secondary-metabolite biosynthesis, oxidoreductase activity and transport) but only 407 genes against FaLH03, while down-regulating secondary-metabolite genes in the conspecific confrontation. Conversely, the *F. avenaceum* isolates showed opposite trends; FaLH03 strongly repressed ribosome-biogenesis and cell-wall genes while inducing oxidative-metabolism pathways, whereas FaveI494 displayed a modest transcriptional response dominated by down-regulation of cell-division and chromosome-segregation genes. Gene-ontology enrichment highlighted an opponent-specific reversal of the *secondary-metabolite biosynthetic process* category in *F. graminearum*: down-regulated in intra-specific confrontation but up-regulated in both inter-specific encounters.

Collectively, our results demonstrate that competitive outcomes are shaped more by strain identity than by species identity, with each strain deploying a distinct molecular arsenal, ranging from metabolite-mediated antagonism to targeted transcriptional shutdown, when confronted with a specific opponent. These findings refine our understanding of *Fusarium* community dynamics and provide a framework for developing strain-targeted biocontrol strategies against FHB.

## Introduction

The *Fusarium* genus comprises over 250 species, most of which are phytopathogenic agents of cereals such as wheat, maize, rice, and barley (Parry et al. 1995; Glenn 2007; Osborne, and Stein 2007). Among them, *Fusarium graminearum* is particularly notable for causing Fusarium head blight (FHB), a devastating plant disease responsible for significant yield losses in wheat and other cereals on a global scale. In addition to its major economic impacts (Windels 2000) FHB represents a serious health concern (Smith, and Thakur 1996; Janik et al. 2020) due to the production of mycotoxins, particularly type B trichothecenes, which persist throughout food processing (Anderson et al. 1989; Bennett, and Klich 2003; Wagacha et al. 2010). These toxins pose substantial risks to human and animal health, making *Fusarium* control a major agricultural challenge.

FHB is caused not by *F. graminearum* alone, but rather by a complex of *Fusarium* (and occasionnally *Microdochium* species) that coexist dynamically in cereal fields. More than 16 different species can be found on a single ear or grain (Xu et al. 2008) interacting to maintain species-specific homeostasis depending on environmental factors and competitive pressures. Recent evidence indicates that competitive rather than synergistic interactions predominate among these co-occurring *Fusarium* species (Dweba et al. 2017; Petrucci et al. 2023) with negative correlations observed between the abundance of different species, suggesting active competition for resources and niche space.

These *Fusarium* species are characterized by their filamentous morphology, consisting of networks of hyphae forming mycelia (McGinnis 1980). The mycelium can take various forms, *e.g.,* cottony, flattened, or aerial, providing survival and adaptability advantages depending on the environment (Takeshita 2016). The morphological diversity of fungi, particularly in terms of mycelial architecture, pigmentation, and sporulation capacity, plays a crucial role in their adaptation and competitive fitness (Liu, and Nizet 2009). Furthermore, morphological changes, particularly in growth patterns and pigmentation, are often correlated with the production of secondary metabolites. These compounds, while not essential for immediate survival, play pivotal roles in development, adaptation, and stress responses (Avalos, and Estrada 2010; Avalos, and Limón 2022). In *Fusarium fujikuroi*, it has been demonstrated that production of fusarubin, a secondary metabolite, is necessary for pigmentation of perithecia, reproductive structures that enable the release of sexual spores (Studt et al. 2012). Secondary metabolites can also serve as chemical communication molecules, participating in quorum sensing phenomena. In *Fusarium*, quorum sensing molecules such as ethyl acetate, 2-methylpropan-1-ol, and 3-methylbutan-1-ol have been described, playing roles in regulating growth and morphogenesis while appearing to induce antifungal effects in response to spore overpopulation (De Clerck et al. 2022). Thus, secondary metabolites are not merely agents of defense or competition, but also play fundamental roles in inter-microbial communication (Tsitsigiannis, and Keller 2007; Zheng et al. 2015) and reflect fungal metabolomic activity in response to environmental stress.

The production of specific secondary metabolites can directly influence competitive outcomes among *Fusarium* species. Enniatins, cyclohexadepsipeptide metabolites produced by *F. avenaceum*, have been shown to inhibit *F. graminearum* development while promoting *F. avenaceum* growth, suggesting a key role in interspecific competition (Ederli et al. 2021). Conversely, *F. graminearum* has been demonstrated to regulate killer toxin genes during competitive interactions with other *Fusarium* species (Petrucci et al. 2025) highlighting the deployment of specialized molecular arsenals during antagonistic encounters. These findings underscore that secondary metabolites serve not only as general stress responses but as targeted competitive weapons tailored to specific fungal opponents.

Within the phyto-pathosystem, numerous fungal species interact with each other, including both inter-specific and intra-specific interactions among *Fusarium* species. These interactions, which can be mutualistic, neutral, or antagonistic, are crucial for plant health and disease outcomes. Interactions can induce morphological and pigmentation changes including variations in mycelial growth, hyphal deformations, and alterations in sporulation patterns. Such modifications may indicate signals of competition or cooperation. The underlying mechanisms remain largely unexplored. Importantly, recent transcriptomic studies have revealed that microbial interactions with *F. graminearum* can trigger important transcriptional reprogramming, suggesting extensive resource reallocation toward specialized competitive functions (Cho et al. 2012; Lee et al. 2014).

Most studies have examined morphology or metabolites in single-strain cultures; few have probed inter-specific interactions at multiple biological scales. Moreover, the extent of intra-specific variation in competitive strategies remains poorly characterized. A comprehensive understanding of how *Fusarium* species and strains interact at morphological, transcriptional, and metabolic levels is essential for elucidating their ecological dynamics and their contributions to FHB epidemics.

In this study, four strains belonging to two co-occurring *Fusarium* species were investigated: *F. graminearum* (strains FgrI349, FgrPH-1) and *F. avenaceum* (strains FaveI494 and FaLH03). Multiple strains of *F. graminearum* and *F. avenaceum* were selected due to previously documented intra-specific genetic diversity in these species (Kulik et al. 2011; Kelly, and Ward 2018), which may translate into distinct competitive strategies. This diversity was initially characterized for each strain in monoculture to establish baseline phenotypic traits and identify differences among strains. Morphological analysis was performed using both macroscopic and microscopic approaches, supplemented by multi-omic strategies including RNA sequencing and non-targeted metabolomics. In the second phase of the study, each strain was confronted with *F. graminearum* FgrI349, chosen as the reference strain, to investigate the morphological and biological consequences of interspecific interactions, which remain poorly understood. Through macroscopic and microscopic observations, we investigated the roles of pigmentation, hyphal deformations (branching patterns, mycelial barriers), and sporulation (conidia, chlamydospores) as indicators of stress or competition in *Fusarium* species. Complementarily, RNA sequencing and non-targeted metabolomics were employed to link morphological traits with underlying transcriptional and metabolic changes. RNA-seq analysis enabled identification of differentially expressed genes and pathways that may drive morphological changes during competition, while metabolomics provided global profiles of compounds modulated during fungal interactions, including mycotoxins, pigments, and other bioactive compounds. We hypothesized that (*i*) strain identity shapes competition outcomes, (*ii*) interspecific encounters induce large-scale transcriptional repression, and (*iii*) *F. graminearum* displays opponent-specific plasticity.

## Materials and methods

### Fungal strains

The strains *F. graminearum* CBS185.32 (or INRA349, Institut Westerdijk, Utrecht, The Netherlands) and the reference PH-1 (FGSC 9075 / NRRL 31084 / PH-1) (Ma et al. 2010; King et al. 2015, 2017), and the strains *F. avenaceum* INRA494 and FaLH03 (Lysøe et al. 2014) were used throughout this study.

### Microscopy

Single cultures (referred to as *monocultures* hereinafter) of each *Fusarium* strains were prepared by inoculating a one millimeter-wide agar plug from working stock cultures at the center of a 3.6 cm Petri dish containing 1 mL of solid Potato Dextrose Agar (PDA, Difco, France) medium. Similarly, co-cultures set as *confrontations* between strains (itself or between *F. graminearum* CBS185.32 and the three other strains) were prepared by inoculating one-millimeter agar plugs from each strain 1.8 cm apart from each other (and 0.9 cm away from the edge of the plates). Cultures were maintained at 25°C in the dark for up to five days. Pictures were taken in bright light using an Oxion Inverso Microscope (Euromex, The Netherlands) and analyzed with Fiji (Schindelin et al. 2012).

### Macroscopic analyses and sample harvest metabolomics and transcriptomics

Monocultures of each strain were prepared by inoculating three millimeter-wide agar plugs at the center of 9 cm-Petri dishes containing 20 mL of solid Potato Dextrose Agar (PDA, Difco, France) medium. Similarly, confrontations were prepared by inoculating three-millimeters agar plugs from each strain 2.2 cm apart from each other (and 3.4 cm away from the edge of the plates). The cultures were maintained at 25°C in the dark for up to six days. Images of samples were taken with a camera (CCD5) at a magnification of ten and an aperture of 1.2. A combination of Fiji and pixel classification followed by k-means segmentation using k = 4 (package *Numpy*, *Python 3.13*) was used to analyze the images, including the measurement of fungal growth area as well as RGB and HSV color channels distribution. Statistical testing was performed on per-image aggregated values (mean pixel values) using six biological replicates with Kruskal-Wallis test (non-parametric ANOVA) followed by Mann-Whitney U pairwise comparison and multiple comparison Benjamini-Hochberg correction (packages *Numpy, Scipy and Statsmodels*, *Python 3.13*). Regarding growth surface analyses, statistical significance was computed with *JASP 0.96* (JASP Team (2026). JASP (Version 0.96.0)[Computer software]), using six biological replicates with Kruskal-Wallis test (non-parametric ANOVA) followed by Dunn’s pairwise comparison and multiple comparison Bonferroni correction. Threshold for significance was set to 0.05 throughout.

Mycelia in confrontation, defined as the zone covering one cm on either side of the interface between the interacting mycelial fronts, was collected for downstream metabolomic and transcriptomic analyses. Both the mycelium and the PDA were collected for metabolomic analyses whereas only the mycelium was scraped off the surface for transcriptomics. Samples were flash-frozen in liquid nitrogen upon harvest and stored at -80°C until further processing.

### Metabolite extraction and metabolomics analysis by UHPLC-HRMS

Metabolite extraction was divided into two successive steps. 5 mL of solvent 1, consisting of an acetonitrile/water/formic acid mixture (ACN/H₂O/FA, 80:19.9:0.1, v/v/v) was added to each sample. Tubes were shaken for 1 hour, then sonicated for 1 hour. After centrifugation (10 min at 14,000 g, 4°C), 2 mL of the supernatant was collected and transferred to a sterile Eppendorf tube. A second extraction was performed with 5 mL of solvent 2 (H₂O/ACN/FA, 80:19.9:0.1, v/v/v) then shaken for 1 hour, sonicated for 1 hour, then centrifuged (10 min at 14,000 g, 4°C). Two millilitres of supernatant were also collected in a second microtube. Finally, 500 µL of each extract (solvent 1 and solvent 2) were mixed in a vial for LC-MS injection. Analyses by ultra-high performance liquid chromatography coupled to high-resolution mass spectrometry (UHPLC-HRMS) were performed using an Ultimate 3000 UHPLC system (Dionex) combined with an LTQ Orbitrap XL mass spectrometer (Thermo Fisher Scientific, Hemel Hempstead, UK). Chromatographic separation was performed on a 2.1 × 150 mm C18 Zorbax Eclipse XDB column with a particle size of 3.5 µm (Agilent Technologies, Santa Clara, CA, USA). The system uses two mobile phases: phase A made up of Milli-Q water spiked with 0.1% formic acid, and phase B made up of acetonitrile acidified to 0.1% formic acid. Chromatographic separation took place at a constant flow rate of 0.3 mL/min, with the following gradient: initially 95% phase B, decreasing to 95% phase A over a period of 10 minutes, followed by a plateau between 10 and 12.5 minutes maintained at 95% phase B. Then, from 12.5 minutes, a 100% B phase was maintained for 1 minute before gradually returning to 95% A phase between 14 and 17 minutes. The temperature of the column was regulated at 40°C, that of the sample carousel at 15°C, and the volume injected was 5 µL. Mass spectrometry detection was carried out using an electrospray ionization (ESI) source, in positive and negative modes, with a resolution of 15,000 (width at half maximum, FWHM, at m/z 400). The mass spectrum was acquired over a range of m/z 100 to 2,000. Spray voltage was set at 3.5 kV and capillary temperature at 300°C. Each full-scale MS acquisition was followed by MS/MS analysis of the three most intense ions, using collision-induced fragmentation (CID) with energy set at 35 arbitrary units. The dynamic exclusion delay was set at 6 seconds. The metabolomic data were analyzed using the peak areas of the metabolites (features). Features with variance <15% and mean intensity values < 10% were filtered before quantile normalization and Pareto-scaling using the MetaboAnalyst software (Pang et al. 2024). Five biological replicates were analyzed by high-resolution mass spectrometry (HR-MS). Graphical representations and further statistical analyses were performed using MetaboAnalyst and R software.

### RNA extraction, library preparation and sequencing

Total RNA was prepared from frozen mycelia by mechanical shearing and tissue disruption using a 5 mm stainless steel bead (IKA steel beads, Thermo Fisher Scientific, Fisher Scientific) and 700 µL of TRIzol® Reagent (Thermo Fisher Scientific, Invitrogen, USA). The mycelia were ground with a TissueLyser® II (Qiagen, Germany) at 2×20 Hz interrupted by a 10 s pause. Then, the homogenate was transferred to a new microcentrifuge tube and they brought at room temperature (15-25°C) for 5 min. We added 140 µL of chloroform to the tube containing the homogenate. After vigorous homogenization samples are incubated at room temperature for 3 minutes. The samples were centrifuged at 12 000 g at 4°C for 15 min and approximately 350 µL of the upper phase (aqueous phase) containing RNA was recovered. This aqueous phase was mixed thoroughly with 1 volumes of freshly prepared 70 % ethanol. Purification was carried out on a RNeasy Mini Spin column (RNeasy Mini kit Qiagen, Qiagen, Germany) according to the manufacturer’s recommendations. A DNase treatment was also carried out according to the manufacturer’s recommendations (ezDNAse I, Thermo Fisher Scientific, Invitrogen, USA). Total RNA was eluted on 32 µL of RNAse-free water. The quantity and quality control of RNA were measured by UV spectrometry (DeNovix, DeNovix Inc, Wilmington, USA) and fluorometric assay (QuantiFluor RNA sample kit, E33110, Promega, USA) and by electrophoresis on 2 % agarose gel (E-gel EX, Invitrogen, USA) with migration performed using E-gel Power Snap (Thermo Fisher Scientific, Invitrogen, USA).

RNA-seq libraries with polyA purification of mRNA were prepared following the manufacturer’s instructions (NEBNext Ultra II RNA Library Prep Kit for Illumina, New England Biolabs, USA) (NEB#E7770-E775S/L) with 300 ng total RNA as biological starting material. Indexing of all our libraries was carried out using the NEBNext Multiplex Oligos for Illumina (Dual Index Primers Set 1) (NEB #7600S/L). Quantification of our libraries is performed using the NEBNext Library Quant Kit (New England Biolabs, USA) (NEB#E7630S/L) following the manufacturer’s instructions. Quality control of the fragment sizes of the libraries was measured on a High sensitivity D1000 ScreenTape with an Agilent 2200 TapeStation system (Agilent Technologies, USA) and processed by Tapestation Analysis software version 5.1. The experiments were performed for three biological replications.

RNA sequencing was performed using the Aviti™ sequencing system (Element Biosciences, San Diego, CA, USA) at the GenoToul platform (INRAE, Toulouse, France). Libraries were prepared according to the manufacturer’s instructions, and sequencing was conducted to generate 50 bp reads.

### RNA-seq data processing, DEG analysis and functional annotation

Sequenced reads were processed with the nf-core pipeline (Pa et al. 2020) rnaseq 3.18.0 (Patel et al. 2024), choosing Salmon as the quantifier tool (Patro et al. 2017). The pipeline was run on the Core Cluster of the Institut Français de Bioinformatique (IFB) (ANR-11-INBS-0013). Differential gene expression analysis was performed using the Limma-Voom package in R (Ritchie et al. 2015). Raw read counts were pre-processed to filter out lowly expressed genes, retaining only those with a minimum count of 5 in at least one sample and a minimum total count of 15 across all samples. For self-confrontations and confrontation conditions, core-gene size factors were computed using only genes present with a minimum count of 20 in all samples, while the default Limma-Voom size factor calculation was applied for monoculture conditions. Differential expression was assessed using linear models for microarray data (Limma), with empirical Bayes moderation to stabilize variance estimates. Multiple testing correction was applied using the Benjamini–Hochberg (BH) method to control the false discovery rate (FDR). Genes were considered differentially expressed (DEG) if they met the criteria of absolute log₂(fold change) ≥ 1 and adjusted p-value (Padj) ≤ 0.05.

Functional annotation and enrichment analysis were performed using the GO.db 3.23.1 package in R/Bioconductor (Gentleman et al. 2004; Huber et al. 2015), assigning Gene Ontology (GO) terms to categorize their biological functions, molecular activities, and involvement in cellular processes. Enrichment analysis was conducted using a hypergeometric test, with multiple testing correction applied *via* the Benjamini–Hochberg (BH) method. Correlation matrices were calculated using Pearson correlation coefficients to assess relationships between gene expression profiles. All graphical representations, including were generated using Python (Matplotlib, Seaborn, Plotly) and R (ggplot2, pheatmap, and base R graphics).

## Results

### Natural phenotypic variability in *F graminearum* FgrI349 and FgrPH-1, and *F. avenaceum* FaveI494 and FaLH03

Strain diversity within *F. graminearum* and *F.* avenaceum leads to marked phenotypic heterogeneity under identical culture conditions, providing the baseline needed to separate stress-induced changes from inherent strain differences. Morphology, growth kinetics and pigmentation of the four strains FgrI349, FgrPH-1, FaveI494, and FaLH03 were quantified (Figure 1). Macroscopically, both *F. graminearum* strains (FgrI349 and FgrPH-1) formed expansive, red-pigmented aerial mycelia (Figure 1A), whereas *F. avenaceum* isolates formed smaller disks, with irregular margins for FaLH03; FaveI494 displayed a flat, beige, continuous mat, while FaLH03 showed a biphasic colony: an older, flat, brown-orange region flanked by a lighter, more aerial, less pigmented peripheral zone. Microscopic observation revealed highly branched, spaced hyphae for the two *F. graminearum* strains FgrI349 and FgrPH-1 (Figure 1B), a dense hyphal network in FaLH03, and long, sparsely branched, vacuolaed hyphae in FaveI494. Radial growth quantification confirmed the superiority of the *F. graminearum* isolates, whose mycelial coverage exceeded 70 % of the Petri-dish surface after five days (mean ± SD = 83.2 % ± 6.9 for FgrI349 and 99.9 % ± 0.001 for FgrPH-1), whereas the *F. avenaceum* strains reached only 32.2 % ± 1.1 (FaveI494) and 28.6 % ± 8.6 (FaLH03) (Figure S1). To assess spatial zonation, each image pixel was characterized based on local texture, color, and intensity, then clustered by kmeans (k = 4) (Figure 1C). All colonies displayed concentric growth zones, primarily distinguished by texture (*e.g.,* fluffy raised hyphae covering the entire surface of FgrPH-1) and pigmentation (*e.g.,* heterogenous color in FaveI494).

**Figure 1.**
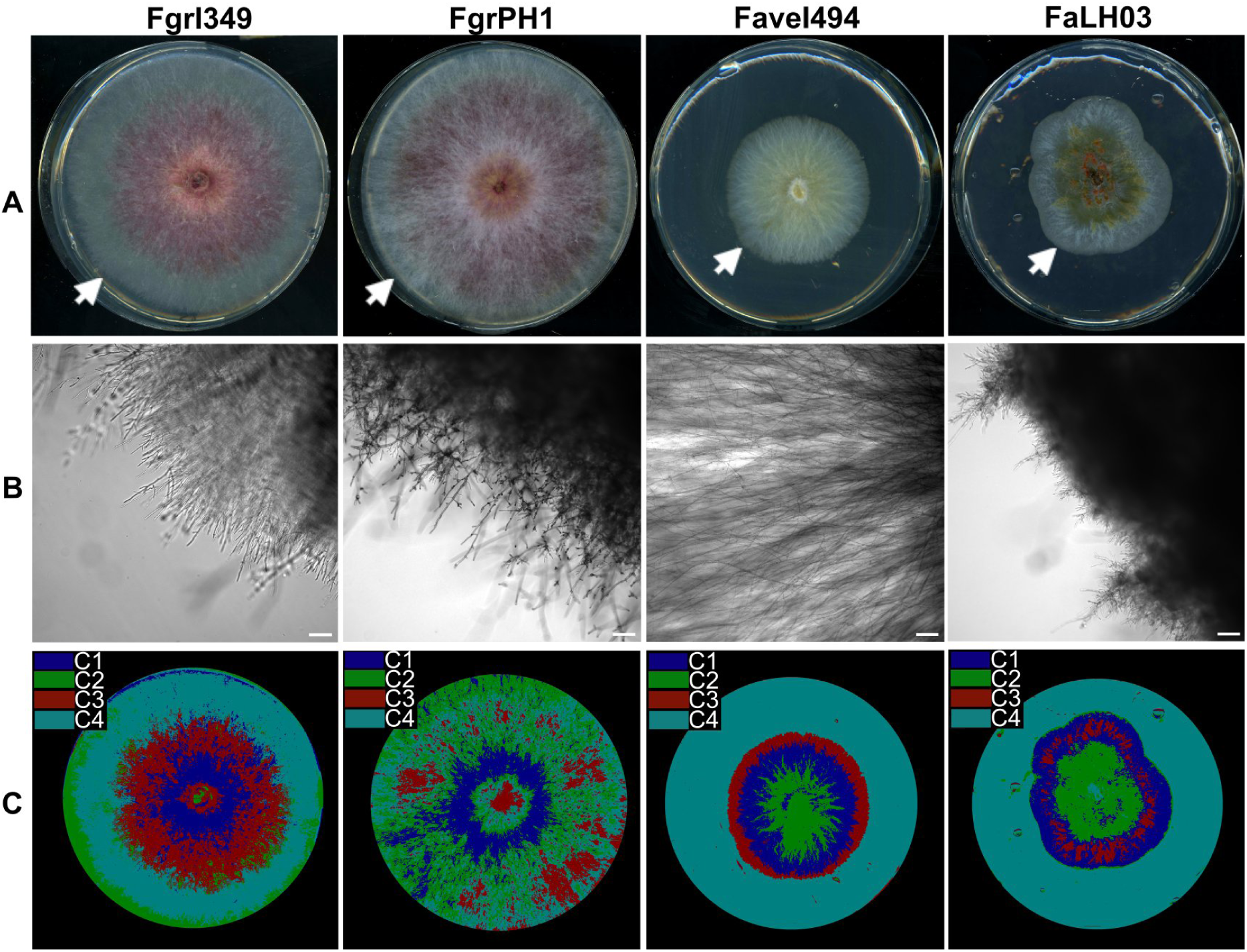
Morphological characteristics of *Fusaria* in monoculture. **A)** FgrI349, FgrPH-1, FaveI494, and FaLH03 grown for five days on PDA, at 25°C in the dark. The white arrows points to growth fronts. **B)** Zoom in (magnification x10) of growth fronts under the microscope. Scale bar = 100 µm. **C)** Visualization of pixel classifications after clustering (k-means with k = 4).

Colorimetric analysis of RGB and HSV channels confirmed these visual differences (Figure 2). The two *F. graminearum* strains showed highly similar red, green and blue distributions, while the *F. avenaceum* strains diverged significantly (Figure 2A-C). FaLH03 exhibited lower red, blue, and green intensities, whereas FaveI494 displayed higher green and blue distributions. Hue values distinguish clearly both *F. avenaceum* strains from *F. graminearum* FgrI349 and FgrPH-1 (Figure 2D). Differences in Hue may reflect variations in the chemical composition of the mycelium (*e.g.,* specific metabolites or enzymes). In Figure 2E, higher Saturation values in *F. avenaceum* strains suggest more vivid coloring, which may reflect denser hyphal networks or increased metabolic activity. Finally, the elevated V values for FaveI494 may indicate differences in hyphal density or surface reflectivity(Figure 2F). Overall, FgrPH-1 and FgrI349 consistently cluster together in growth, texture and colour, whereas, FaLH03 frequently stands out, indicating heterogenous metabolic and morphological phenotypes within *F. avenaceum*.

**Figure 2.**
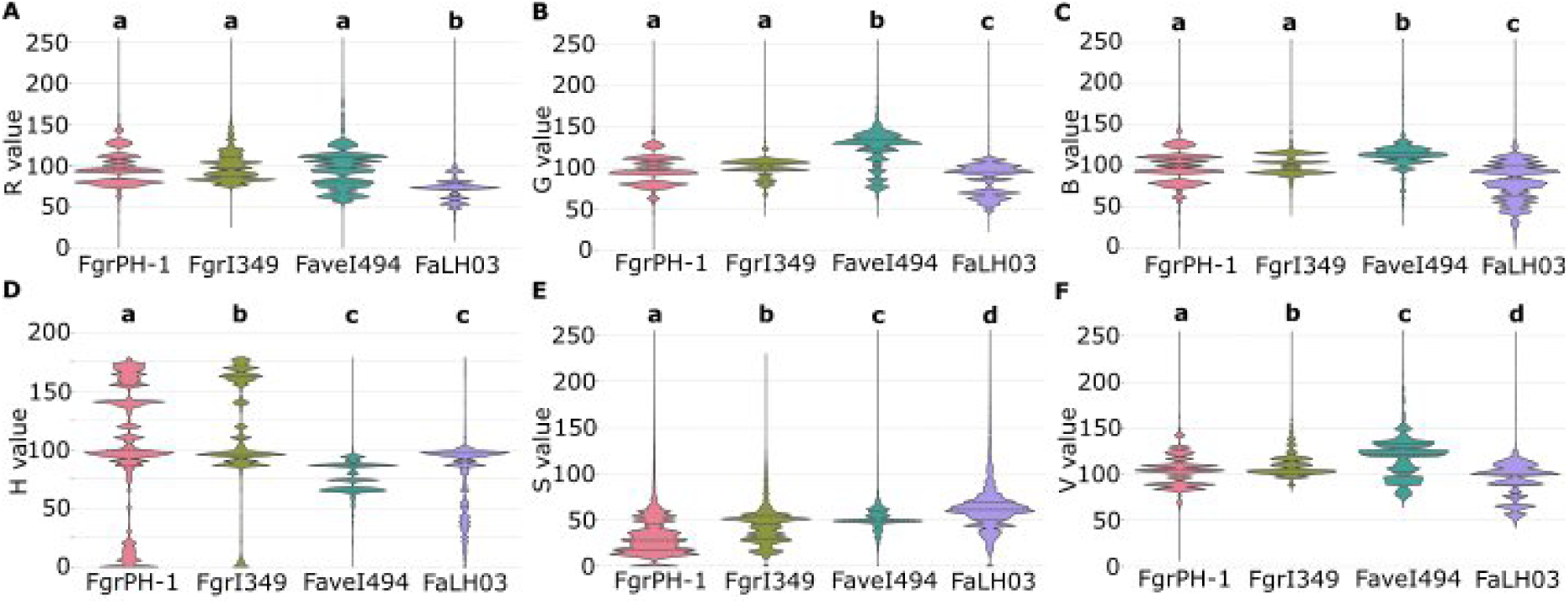
Comparison of RGB and HSV distributions across the four fungal strains. **A)** Distribution of Red channel intensity. **B)** Distribution of Green channel intensity. **C)** Distribution of blue channel intensity. **D)** Distribution of Hue, reflecting color type. **E)** Distribution of Saturation, reflecting color intensity. **F)** Distribution of Value, reflecting brightness. The letters a,b,c, and d indicate the statistical groupings based on Kruskal-Wallis test (non-parametric ANOVA) on per-image aggregated mean pixel values (n = 6) followed by Mann-Whitney U pairwise comparison and multiple comparison Benjamini-Hochberg correction results (p < 0.05).

### Metabolic profiling of FgrI349, FgrPH-1, *Fave*I494, and FaLH03

Untargeted LC-HRMS analyses were carried out to characterize the metabolomic profiles of FgrI349, FgrPH-1, FaveI494, and FaLH03. After metabolomic data acquisition and filtering, a total of 1,008 features were retained for further analysis. Replicate-wise correlation confirmed that each strain formed a distinct cluster, with the two F. graminearum isolates showing tight inter-strain correlation (Figure S2). In contrast, FaLH03 behaves as an outlier, displaying no appreciable correlation with either FaveI494 or the *F. graminearum* strains, underscoring the pronounced intra-species heterogeneity observed phenotypically.

The upset plot in Figure 3A illustrates the feature distribution between strains. Individual strain counts ranged from 471 (FgrI349) to 555 (FgrPH-1). Only 23 features were shared by all four strains, whereas 115 features were specifically found in both *F. graminearum* isolates (out of the 431 features non-exclusively shared between both strains) and merely 49 between the two *F. avenaceum* isolates (out of the 115 features non-exclusively shared between FaveI494 and FaLH03). The largest exclusive set comprised 295 features (≈ 61 % of FaLH03’s metabolites), whereas FaveI494, FgrI349 and FgrPH-1 possessed merely 69, 12 and 1 strain-specific features, respectively, highlighting the relatively homogeneous metabolome of *F. graminearum* versus the diversified chemistry of *F. avenaceum*.

**Figure 3.**
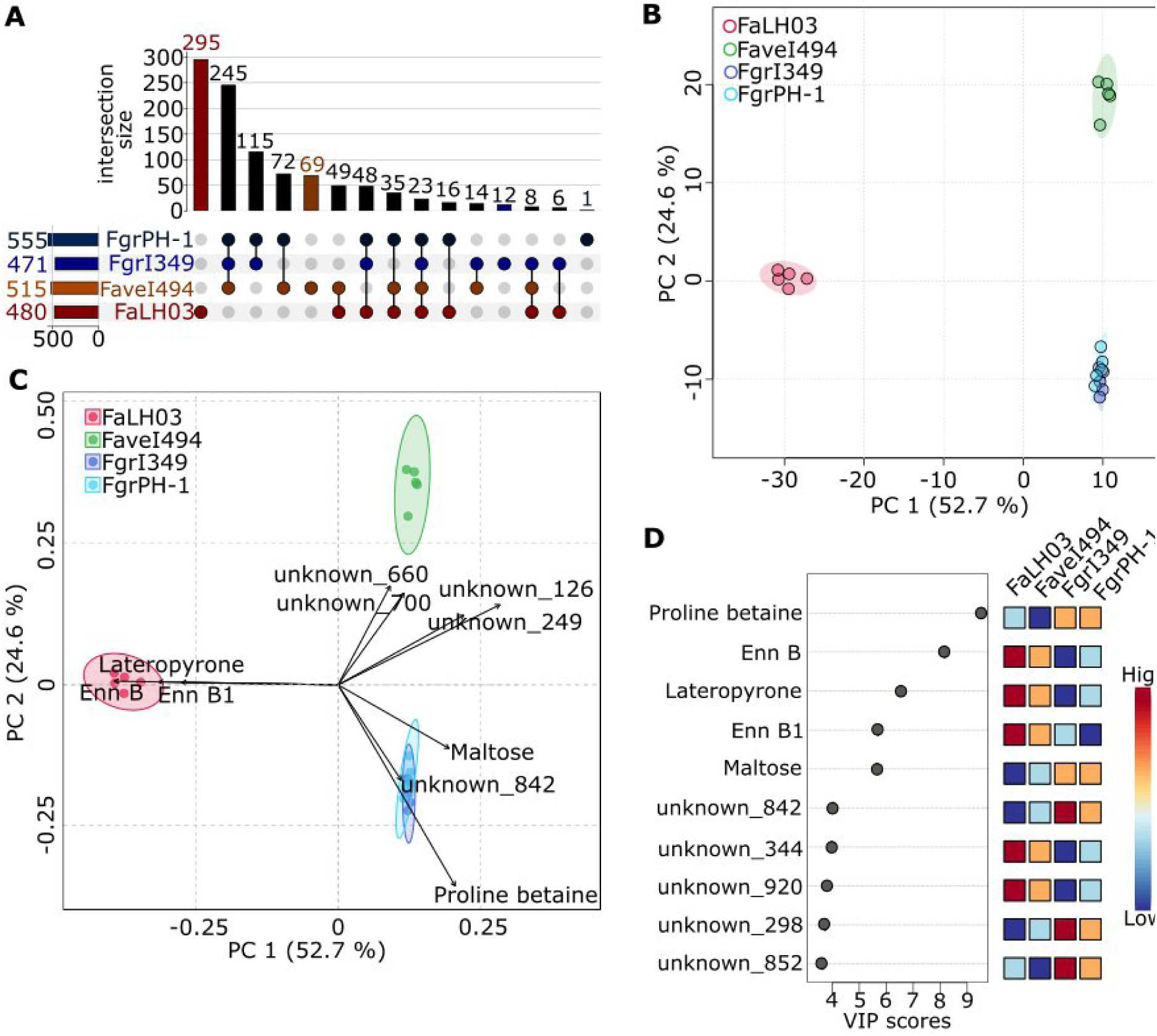
Metabolomic profiling of *F. avenaceum* FaLH03 and FaveI494, and *F. graminearum* FgrI349 and FgrPH-1, grown in monocultures. **A)** Feature distribution across strains (upset plot). **B)** Principal component analysis (PCA) scores plot of metabolomic profiles. The ellipses around each group represent the 95% confidence interval for the distribution of samples within that group. **C)** PCA biplot showing the top 10 (magnitude) loading vectors **D)** Variable Importance in Projection (VIP) score plot within the context of Partial Least Squares (PLS) regression of the metabolites, featuring the top 10 features. The heatmap on the right shows the relative abundance of each metabolite feature across FaLH03, FaveI494, FgrI349, and FgrPH-1. The color scale ranges from blue (low abundance) to red (high abundance).

Principal-component analysis captured 77.3 % of the variance over the first two dimensions (PC1 = 52.7 %, PC2 = 24.6 %) (Figure 3B). PC1 cleanly separated FaLH03 (red) from the remaining strains, which clustered on the opposite side. Within that cluster, the two *F. graminearum* strains overlapped closely, while FaveI494 (green) was distinguished along PC2. The loadings plots identified a handful of discriminating features (Figure S3 and Figure 3C). The putative features enniatin B, enniatin B1 and lateropyrone drove the FaLH03 position, maltose, proline betaine and unknown_842 anchored the *F. graminearum* group, and unknown_660, unknown_700, unknown_126 and unknown_249 characterized FaveI494. Partial-least-squares VIP analysis (Figure 3D) confirmed the same species-level split. Five of the top-10 VIP metabolites were annotated: proline betaine and maltose (present in both *F. graminearum* strains) and enniatin B, enniatin B1 and lateropyrone (enriched in the *F. avenaceum* strains). The remaining high-VIP features remained unassigned but contributed markedly to the separation. Collectively, the metabolomic data reveal stark chemical divergence between the two species and, within *F. avenaceum*, a pronounced strain-specific metabolic fingerprint contrasted with the relative uniformity of *F. graminearum*. To elucidate the transcriptional programs underlying these metabolic patterns, we next performed RNA-seq on the same set of cultures.

### Transcriptomic profiling of FgrI349, FgrPH-1, *Fave*I494, and FaLH03

Transcriptomic profiling of FgrI349, FgrPH-1, FaveI494, and FaLH03 grown in monocultures revealed markedly different regulatory landscapes. Global expression profiles of the two *F. graminearum* isolates were highly concordant (pairwise Pearson correlation r = 0.89–0.94 across replicates; Figure 4A, left panel), whereas the *F. avenaceum* strains displayed only modest concordance (r = 0.38-0.53; Figure 4A, right panel), reinforcing the pronounced intra-species variability suggested by morphology and metabolomics analyses. Pairwise differential-expression analysis identified 1,632 up-and 1,074 down-regulated genes in FgrPH-1 relative to FgrI349; Figure 4B, left panel), whereas a substantially larger set of genes varied between FaveI494 and FaLH03, with 2,682 up-and 3,943 down-regulated genes in FaveI494 relative to FaLH03 (Figure 4B, right panel). Thus, the *F. graminearum* strains share a comparatively conserved transcriptional program, whereas the *F. avenaceum* isolates exhibit substantially greater regulatory divergence that may underpin their distinct phenotypes.

**Figure 4.**
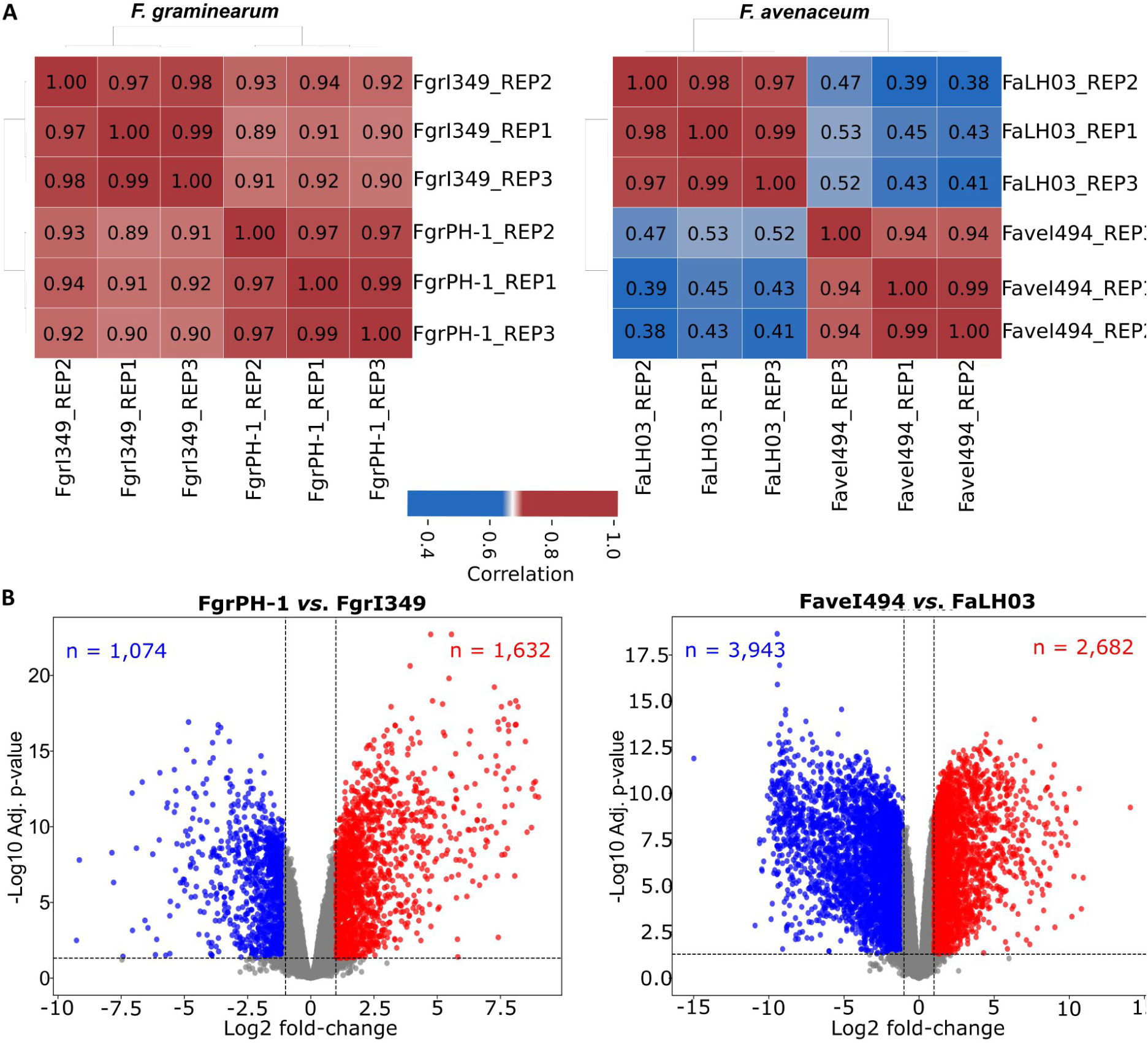
Transcriptomic profiling of *F. graminearum* FgrPH-1 and FgrI349, and *F. avenaceum* FaveI494 and FaLH03. **(A)** Correlation heatmaps of transcriptomic profiles between *F. graminearum* strains (left) and *F. avenaceum* strains (right). The color scale ranges from blue (low correlation, 0.4) to red (high correlation, 1.0). **(B)** Volcano plots of differential gene expression between *F. graminearum* strains FgrPH-1 *vs*. FgrI349 (left), and *F. avenaceum* strains FaveI494 *vs*. FaLH03 (right).

Gene-ontology enrichment of the *F. graminearum* differentially expressed genes (Figure S4A) highlighted up-regulation of secondary metabolite biosynthesis together with monooxygenase, oxidoreductase, and transmembrane transporter activities (and associated heme/iron-binding functions), indicating a metabolically versatile, growth-supporting state in FgrPH-1 (Eagan, and Keller 2025). Down-regulated genes were enriched for nucleolus, rRNA processing, ribosome biogenesis, and related small-subunit processome/preribosome categories, suggesting that FgrPH-1 may achieve rapid growth by allocating fewer transcriptional resources to ribosome production while maintaining efficient translation. This interpretation aligns with a model in which enhanced metabolic efficiency and optimized energy utilization contribute to the strain’s improved growth phenotype (Shore, and Albert 2022). In *F. avenaceum*, GO analysis (Figure S4B) showed that down-regulated genes in FaveI494 were strongly enriched for core metabolic categories, such as small molecule, carboxylic acid, oxoacid, and amino acid metabolic processes, together with catalytic and oxidoreductase activities, detoxification, and response to toxic substance, pointing to a broad reduction in primary metabolic and redox/detoxification activity relative to FaLH03. Conversely, up-regulated genes clustered in nuclear-and organelle-associated components (nuclear lumen, nucleus, nucleolus, intracellular organelle lumen, membrane-enclosed lumen) along with chromosome organization and pre-ribosome assembly, suggesting a transcriptional shift toward nuclear and ribosomal biogenesis activity rather than a wholesale metabolic overhaul.

Having established the baseline of morphological, metabolomic and transcriptomic signatures of FgrI349, FgrPH-1, FaveI494 and FaLH03 in monoculture, we next sought to explore how these traits evolve during intra-and inter-specific confrontations, pairing each strain against FgrI349 or against itself, to uncover the dynamic molecular responses that emerge when the fungi coexist.

### Morphological and metabolic profiling of FgrPH-1, FaveI494, and FaLH03 with FgrI349 grown in self-confrontation co-cultures

To determine whether identical genotypes influence each other’s phenotype, we performed self-confrontation (SC) assays, *i.e.,* each strain *versus* itself (Figure 5). Macroscopic views (left and center panels) display colony outlines and the encounter zone, while the inset microscopy (right panels) shows hyphal interdigitation that is indistinguishable from monoculture morphology. Ranking of colony expansion reproduced the monoculture hierarchy (SC-FgrPH-1 > SC-FgrI349 > SC-FaveI494 > SC-FaLH03; Figure S5A), indicating that proximity to an identical partner does not remarkably alter growth.

**Figure 5.**
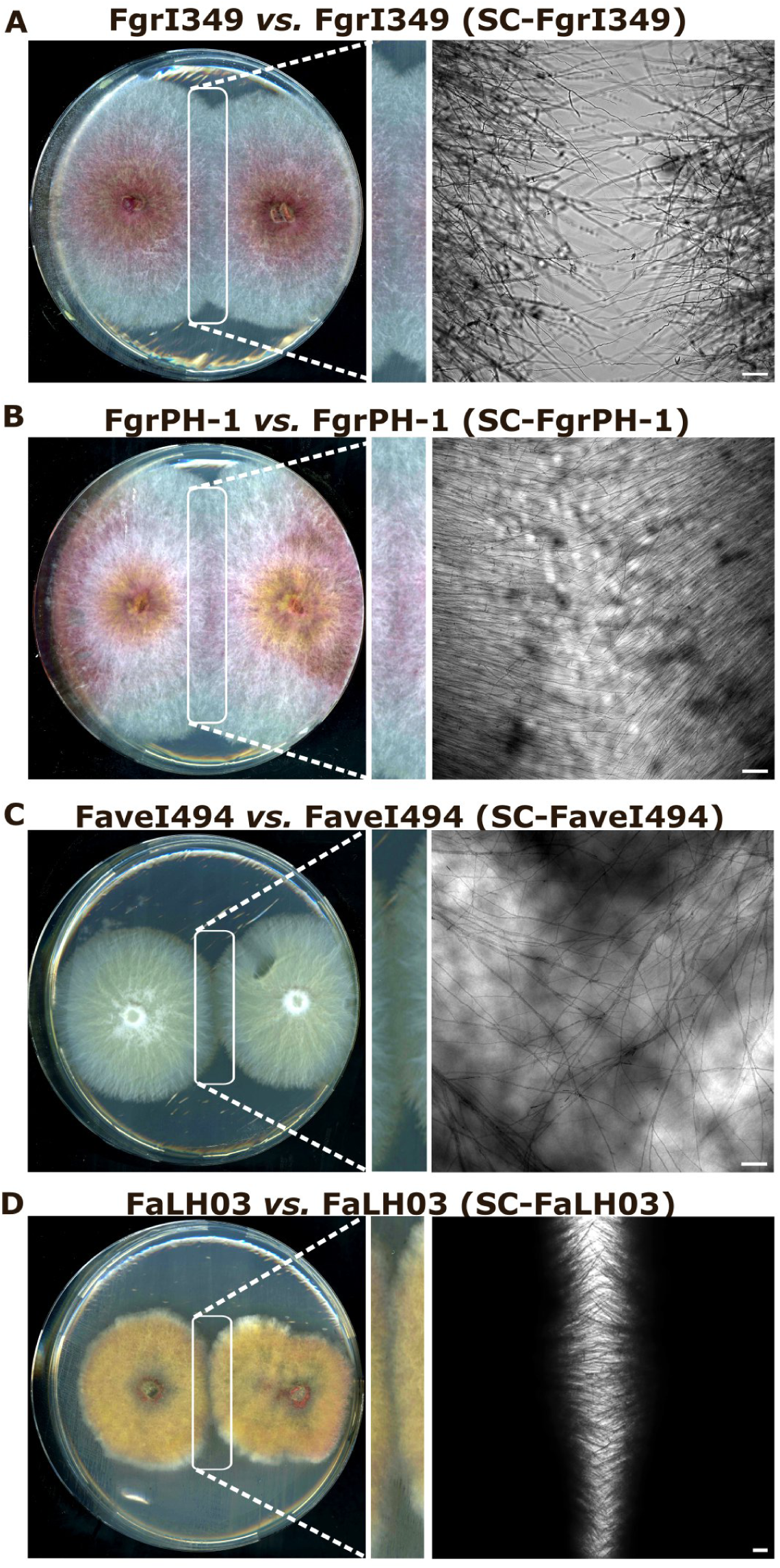
Morphological characteristics of FgrI349 **(A)**, FgrPH-1 **(B)**, FaveI494 **(C)**, and FaLH03 **(D)** in self-confrontations (SC), grown for four days on PDA, at 25°C in the dark. For each self-confrontation, the left image shows the macroscopic view of the fungal colonies on agar plates, with a rectangular inset indicating the region magnified on the right. The right image provides a microscopic view of the confrontation zone, highlighting the hyphal interactions. The scale bar indicates 100 µm.

Untargeted metabolomics of the SC zones yielded 977 features. Replicate correlation confirmed that each SC profile clustered tightly with its corresponding monoculture, similar to the strain-specific patterns reported previously (Figure S5B). FaLH03 again behaved as an outlier, showing no significant correlation with either SC-FaveI494 or any *F. graminearum* profiles. The UpSet plot in Figure S6A visualizes feature intersections across SC and monoculture conditions. The largest shared block (n = 229) comprises metabolites common to SC-FaLH03 and its monoculture, whereas a second block (n = 182) contains features shared by all strains and conditions except those involving FaLH03, further emphasizing the distinct metabolic network of this strain. PCA of the combined monoculture and SC datasets (PC1 = 48.8 %, PC2 = 20.5 %; 77.3 % total variance) placed each SC sample directly onto its monoculture counterpart (Figure S6B). Loadings (Figure S6C) identified the same discriminating metabolites as in the monoculture analysis: enniatin B and lateropyrone remained tightly linked to FaLH03, maltose, proline betaine and unknown_842 to the *F. graminearum* strains, and unknown_660, unknown_197, unknown_126 and unknown_249 to FaveI494.

Transcriptomic divergence between conspecific strains under a controlled, non-antagonistic interaction context, was characterized by comparing self-confrontation (SC) transcriptomes within each species. Within *F. graminearum*, self-confronted FgrPH-1 and FgrI349 differed at 2,106 genes (828 down-and 1,278 up-regulated in FgrPH-1 relative to FgrI349; Figure 6A), a magnitude comparable to the FgrPH-1 *vs.* FgrI349 monoculture comparison reported earlier (2,706 genes). Within *F. avenaceum*, FaveI494 and FaLH03 diverged far more substantially under self-confrontation, with 6,381 genes differentially expressed (3,717 down-and 2,664 up-regulated in FaveI494 relative to FaLH03; Figure 6B), *i.e.,* substantially more genes than either *F. graminearum* strain pair (2.4-fold relative to the monoculture comparison; 3.0-fold relative to the self-confrontation comparison). This is consistent with the substantially lower inter-strain correlation observed for *F. avenaceum* isolates in the correlation analysis (Figure 4B) and reinforces the pattern of pronounced intra-species transcriptional variability in *F. avenaceum* relative to *F. graminearum*, evident even under an interaction-matched (self-confrontation) design.

**Figure 6.**
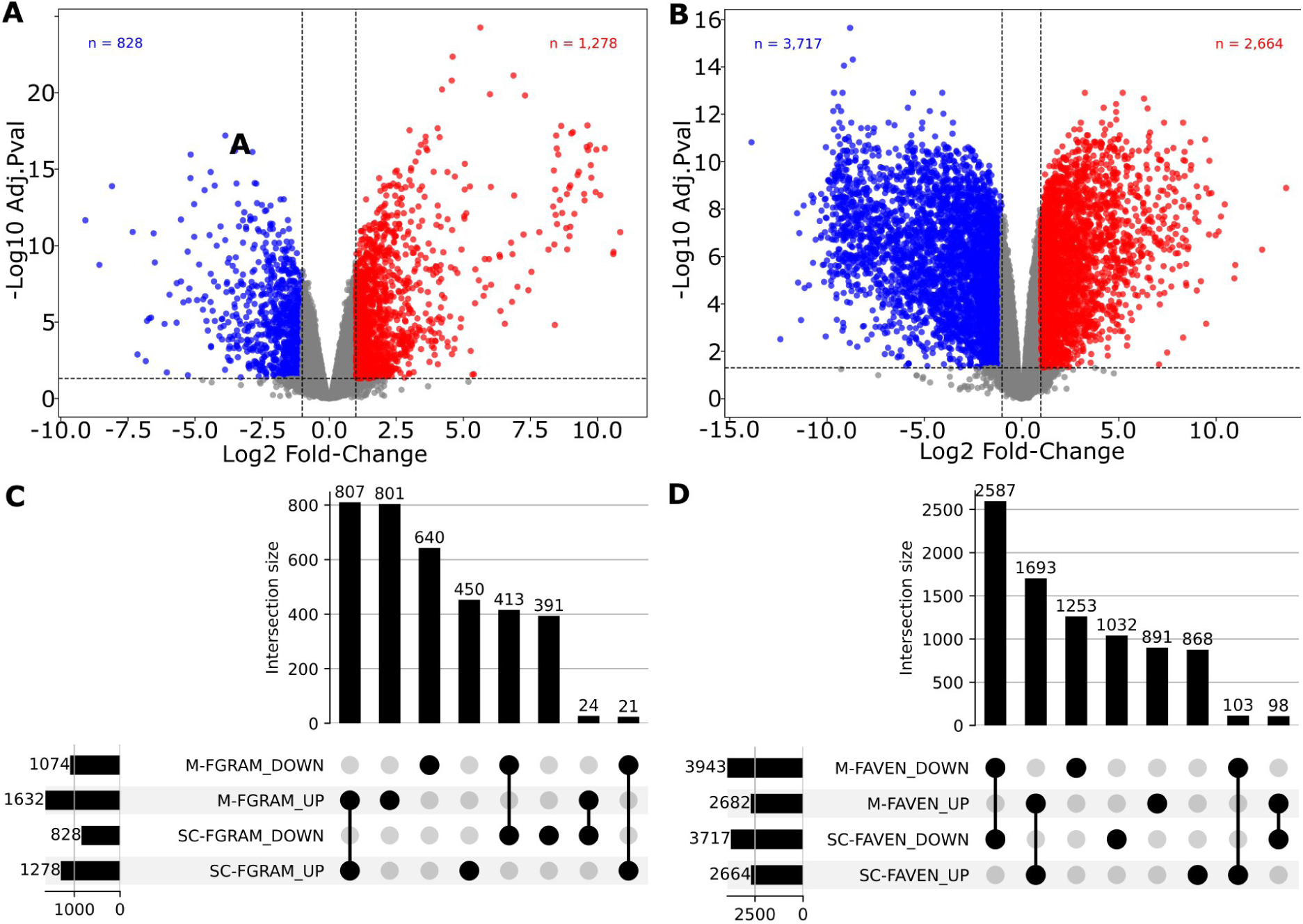
Transcriptomic characteristics of self-confrontation (SC) cultures. **(A)** Volcano plot of differential gene expression between *F. graminearum* strains FgrPH-1 and FgrI349 in self-confrontation (n = 828 down-, n = 1,278 up-regulated genes in FgrPH-1 relative to FgrI349). **(B)** Volcano plot of differential gene expression between F. avenaceum strains FaveI494 and FaLH03 in self-confrontation (n = 3,717 down-, n = 2,664 up-regulated genes in FaveI494 relative to FaLH03). **(C)** and **(D)** UpSet plots comparing differentially expressed gene sets identified in monoculture (M) *versus* self-confrontation (SC) for the same strain pairs, in *F. graminearum* **(C)** and *F. avenaceum* **(D)**. Horizontal bars indicate the total number of genes in each set; vertical bars indicate the number of genes in each intersection, shown for the top eight intersections.

GO enrichment of the FgrPH-1 *vs.* FgrI349 self-confrontation DEGs (Figure S7A) showed genes up-regulated in FgrPH-1 enriched for secondary metabolite biosynthetic process, transmembrane transport, and transmembrane transporter activity, while genes lower in FgrPH-1 (*i.e.,* up in FgrI349) were enriched for oxidoreductase activity and membrane-associated components; secondary metabolite biosynthetic process was notably enriched on both sides, indicating that secondary metabolism differs qualitatively rather than uniformly between the two strains, with distinct gene subsets favored in each. GO enrichment of the FaveI494 *vs.* FaLH03 self-confrontation DEGs (Figure S7B) revealed a sharp and coherent pattern. Genes up-regulated in FaveI494 were dominated by ribosome-and nucleolus-associated categories (nucleolus, preribosome, rRNA processing, rRNA metabolic process, and ribosome biogenesis) while genes lower in FaveI494 (up in FaLH03) were enriched for oxidoreductase activity, cellular response to oxidative stress, and general small-molecule/carboxylic acid metabolic processes. To assess whether these strain-pair differences reflect constitutive divergence or are specific to the self-confrontation context, we compared the SC-derived DEG sets against the corresponding monoculture comparison of the same strain pairs. For *F. graminearum* (Figure 6C), 413 of 1,074 M-down (monocultures) genes (38 %) were also down under SC, and 807 of 1,632 M-up genes (49 %) were also up under SC, with only 45 genes (∼1 % of the total DEG union) showing discordant direction between conditions. Condition-specific up-and down-regulated genes represented 49 % (801 genes of 1,632) and 60 % (640 genes of 1,074) in monoculture comparisons, and 35 % (450 genes of 1,278) and 47 % (391 genes of 828) self-confrontation comparisons. For *F. avenaceum*, concordance between M and SC was high (Figure 6D): 2,587 of 3,943 M-down genes (66 %) were also down under SC, and 1,693 of 2,682 M-up genes (63%) were also up under SC, with only 201 genes (∼3 % of the total DEG union) showing discordant direction between conditions.

Together, our results indicate that self-recognition does not provoke major reprogramming in either species under our conditions. Having established this baseline, we next examined non-self (intra-and inter-specific) confrontations between FgrI349 and each of the other strains to uncover the biochemical and phenotypic shifts associated with genuine competitive interactions.

### Macro-and microscopic observations of inter-and intra-specific confrontations of FgrPH-1, FaveI494, or FaLH03 *vs.* FgrI349

We confronted the reference strain FgrI349 with its con-specific FgrPH-1 (C-FgrPH-1) and with the two *F. avenaceum* isolates FaveI494 (C-FaveI494) and FaLH03 (C-FaLH03). Macroscopic photographs (Figure 7, left panels) display colony outlines and the interface zones, while the rectangular insets and the corresponding microscopic panels reveal hyphal organization within those zones. In the intra-specific pairing C-FgrPH-1 (Figure 7A), a faint boundary formed at the contact line, the mycelium appearing less dense. The microscope field shows a network of septate, branching hyphae (with an estimated diameter of 3-4 µm) crossing at multiple angles from different directions, consistent with two colony fronts of the same species merging into a shared mycelial mat, without obvious antagonism, which suggests a neutral interaction. By contrast, the inter-specific encounter with FaveI494 (C-FaveI494, Figure 7B) produced a conspicuous delimitation marked by intense red pigmentation (left panel and insert). The brightfield observation of this zone reveals that *F. graminearum* FgrI349 displays a dense, irregularly cross-hatched hyphal mesh (estimated diameter of 4 µm) crossing at many angles, while *F. avenaceum* FaveI494 shows sparser, narrower (estimated diameter < 3 µm), strongly parallel-aligned hyphae. In the color close-up of FaveI494 mycelium (right panel), individual hyphae appear as distinctly darker, more saturated carmine-red hyphal strands (white arrows) against a paler pink background, indicating the concentration of a red pigment

**Figure 7.**
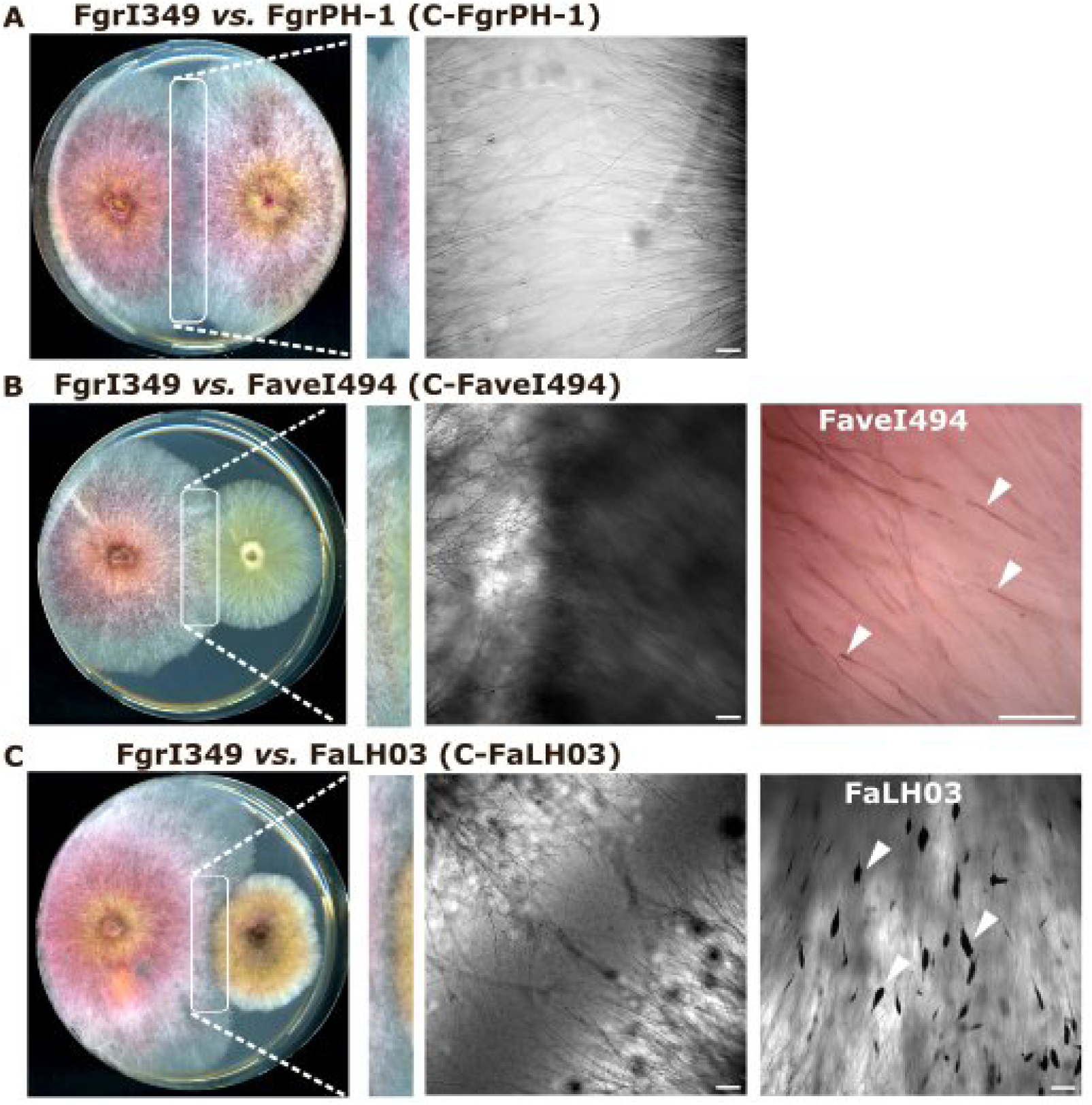
Morphological characteristics of FgrPH-1 **(A)**, FaveI494 **(B)**, and FaLH03 **(C)** in confrontations with FgrI349, grown for four days on PDA, at 25°C in the dark. For each confrontation, the left image shows the macroscopic view of the fungal colonies on agar plates, with a rectangular inset indicating the region magnified on the right. The images on the right provide microscopic views of the confrontation zone. Scale bar = 100 µm.

in the fungal biomass rather than being a uniform stain across the whole field. The confrontation with FaLH03 (C-FaLH03) differed further (Figure 7C), the two colonies remaining physically separated by an unbridged gap. The FgrI349 side shows sparser, thinner, more randomly branching hyphae than the FaLH03 side, which shows a dense fan of aligned hyphae studded with numerous dark blobs. Focusing on the FaLH03 region, numerous discrete fusiform to ovoid dark bodies (approximate measurements 60–150 µm long × 15–65 µm wide) can be observed, oriented along the local hyphal growth axis. Together, these three confrontations reveal a gradient of interaction phenotypes. The two *F. avenaceum* isolates particularly diverged from one another in their specific response, further highlighting isolate-level variation in competitive strategy.

Growth area quantification revealed that FgrI349 expanded significantly more when facing either *F. avenaceum* strain than in its self-confrontation, whereas its growth was unchanged in the intra-specific duel with FgrPH-1 (Figure 8A). Conversely, FgrPH-1 achieved a larger occupied area when paired with FgrI349. Regarding *F. avenaceum*, FaveI494 showed reduced expansion in its heterologous confrontation, and FaLH03 displayed no significant deviation from its self-confrontation (Figure 8C). Colorimetric analysis of the confrontation zones highlighted strain-specific shifts in RGB and HSV channels. In the intra-specific case, FgrI349 exhibited higher red intensity and increased values for hue, saturation and brightness compared with its self-confrontation, whereas only saturation rose for FgrPH-1 (Figure S8). Inter-specific interactions elicited broader changes: FgrI349 showed significant elevation of all RGB channels and the HSV value component against both *F. avenaceum* strains, which may indicate heightened metabolic activity (Figure S9). FaLH03 mirrored this increase, while FaveI494 displayed a modest decline in RGB and value measures, suggesting stress or resource limitation. Hue responses diverged. FgrI349’s hue increased when confronting FaLH03 but decreased against FaveI494, while saturation varied in a strain-dependent manner. These color shifts, quantified by RGB/HSV parameters, likely reflect underlying metabolic and physiological reprogramming that accompanies the distinct morphological outcomes observed during inter-specific competition.

**Figure 8.**
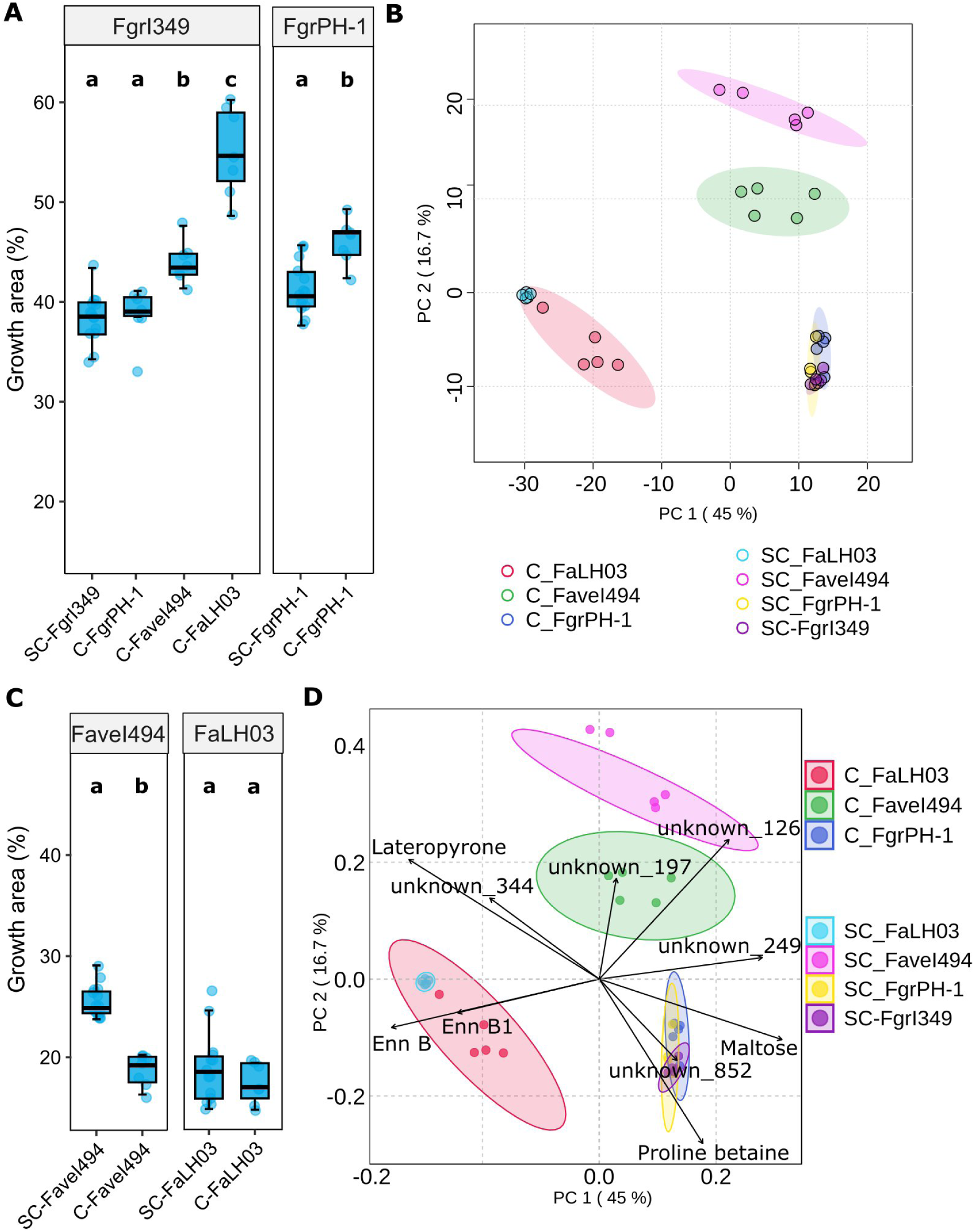
Characteristics of confrontation *vs.* self-confrontation cultures. **(A)** and **(C)** Fungal growth measured as the surface of the Petri dish occupied by mycelium, in percentage of the total area, in *F. graminearum* FgrI349 and FgrPH-1 **(A)**, and *F. avenaceum* FaveI494 and FaLH03 **(C)**. The letters a, b, and c indicate statistical groupings based on Kruskal-Wallis test (non-parametric ANOVA) followed by Dunn’s post hoc (p < 0.05). **(B)** and **(D)** Principal Component Analysis (PCA) showing the distribution of samples from seven groups: C_FaLH03 (red), C_FaveI494 (green), C_FgrPH-1 (blue), SC_FaLH03 (cyan), SC_FaveI494 (pink), SC_FgrPH-1 (yellow), and SC_FgrI349 (purple). Each ellipse represents the 95% confidence interval for the corresponding group. **(B)** shows the scores plot and **(D)** the biplot, with arrows indicating the loading vectors of variables contributing most to the separation among groups.

### Metabolic profiling of FgrI349 in confrontation with FgrPH-1, FaveI494, and FaLH03

Untargeted LC-HRMS of the confrontation zones for C-FgrPH-1, C-FaveI494, C-FaLH03 together with the corresponding self-confrontations (SC) yielded 938 reliable features. Replicate-wise Pearson correlation confirmed tight clustering of each condition and, notably, strong concordance between each heterologous confrontation and the matching SC profile of the non-*F. graminearum* partner (Figure S10). All samples involving only *F. graminearum* isolates (SC-FgrI349, SC-FgrPH-1, C-FgrPH-1) also displayed high mutual correlation, supporting the view that strain-specific inter-specific interactions drive the observed metabolic distinctions. PCA of the seven sample groups captured 61.7 % of the total variance (PC1 = 45 %, PC2 = 16.7 %). The scores plot (Figure 8B) resolved four clusters: a blue cluster comprising the *F. graminearum-*only cultures (SC-FgrI349, SC-FgrPH-1, C-FgrPH-1); a red cluster containing SC-FaLH03 and C-FaLH03, indicating that the FaLH03 metabolome dominates any interaction in which it participates; a pink cluster for SC-FaveI494; and a green cluster for C-FaveI494, whose separation reflects the influence of FgrI349 on the metabolic milieu. Loadings (Figure S11) showed that most variables occupy a dense central region, with only a handful of metabolites contributing to the dispersion. The PCA biplot (Figure 8D) highlighted the same ten dominant features as in monoculture: enniatin B, enniatin B1 and lateropyrone (together with unknown_344) aligned with FaLH03, whereas maltose, proline betaine and unknown_852 tracked the *F. graminearum* samples. Hierarchical clustering of the 50 most variable features (Figure S12) revealed distinct abundance patterns across the seven conditions. A first cluster of 20 features (unknown_974–unknown_1027) was enriched in all FaLH03–containing cultures (SC-FaLH03 and C-FaLH03), suggesting FaLH03-specific signature metabolites. A second cluster of 12 features (unknown_775–unknown_1156) rose exclusively in SC-FaLH03, hinting at a unique self-response. A third cluster of four metabolites (unknown_1165, unknown_914, unknown_178, unknown_175) characterized *F. graminearum* intra-species cultures irrespective of strain, while a fourth cluster of 14 features was most abundant in FaveI494, particularly under self-confrontation. Together, these patterns confirm that each *F. avenaceum* strain possesses a characteristic metabolic fingerprint that persists across self-and cross-species contacts, whereas *F. graminearum* strains maintain a comparatively homogeneous metabolomic profile Having established that confrontation-induced metabolic reprogramming differs between the species, we next interrogated whether parallel transcriptional changes accompany these chemical shifts by performing RNA-seq on the same set of cultures.

### Transcriptional dynamics during inter-and intra-specific confrontations with *F. graminearum* FgrI349

To uncover the molecular pathways engaged during self-recognition and heterologous competition, we performed RNA-seq on mycelia collected at the confrontation zone of self-confrontations (SC-FgrI349, SC-FgrPH-1, SC-FaveI494) and confrontations of each partner with FgrI349 (C-FgrPH-1, C-FaveI494, C-FaLH03). Because *F. graminearum* and *F. avenaceum* possess distinct gene repertoires, we analyzed the two species separately throughout. In addition, the C-FgrPH-1 condition could not distinguish transcripts originating from FgrI349 from those originating from FgrPH-1.

Global transcriptional profiles were first examined by PCA, starting with *F. graminearum* (Figure 9A). PCA of FgrI349 in conspecific confrontation with FgrPH-1 and interspecific confrontations with FaveI494 and FaLH03, as well as self-confrontation samples for FgrI349 and FgrPH-1, resolved five distinct clusters, one per condition (PC1 = 42.3%, PC2 = 16.8%; 59.1% of total variance). Both interspecific confrontations (C-FaveI494 and C-FaLH03), representing FgrI349’s transcriptional signal when facing each *F. avenaceum* partner, separated clearly from FgrI349’s own self-confrontation baseline (SC-FgrI349) along PC1 and PC2, confirming that FgrI349 mounts a genuine, partner-associated transcriptional shift rather than remaining transcriptionally static. Notably, the magnitude of this shift differed by opponent: C-FaveI494 was markedly further from SC-FgrI349 than C-FaLH03 was, indicating that FgrI349’s response is substantially stronger when confronting FaveI494 than when confronting FaLH03. The intra-specific pairing showed a parallel pattern, with C-FgrPH-1 (the necessarily blended FgrPH-1/FgrI349 signal) separating from SC-FgrPH-1 along both PC1 and PC2, suggesting the mixed confrontation transcriptome likewise diverges from either parental self-confrontation baseline rather than reading as an intermediate of the two.

**Figure 9.**
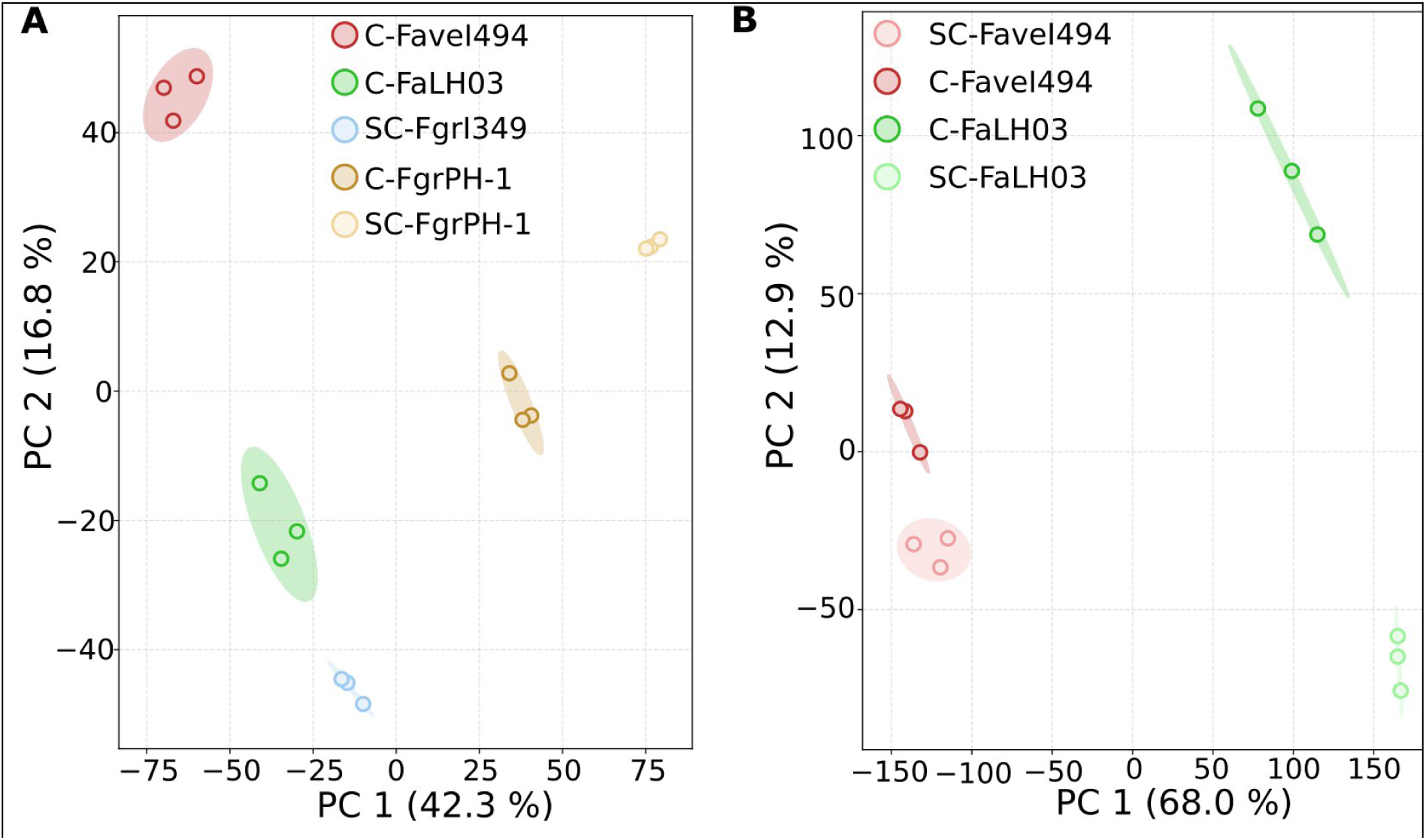
Principal Component Analysis (PCA) of gene expression profiles measured in *Fusarium* in confrontations. **(A)** PCA scores plot showing the separation of SC-FgrI349, SC-FgrPH-1, C-FgrPH-1, C-FaveI494, and C-FaLH03 based on their *F. graminearum* gene expression profiles. **(B)** PCA scores plot showing the separation of SC-FaLH03, SC-FaveI494, C-FaLH03, and C-FaveI494 based on their *F. avenaceum* gene expression profiles. Each point represents an individual sample (n=3 per group), and colored ellipses denote 95 % confidence intervals.

For *F. avenaceum* (Figure 9B), PCA of the two strains’ confrontation and self-confrontation samples (PC1 = 68.0%, PC2 = 12.9%; 80.9% of total variance) revealed a striking asymmetry between the two isolates. FaveI494’s confrontation and self-confrontation samples (C-FaveI494, SC-FaveI494) remained close together, occupying adjacent, only modestly separated positions along PC1, indicating that FaveI494’s transcriptome is comparatively stable across the two conditions. FaLH03, by contrast, showed a dramatic separation between C-FaLH03 and SC-FaLH03 spanning both PC1 and PC2, with confrontation samples displaced far from their own self-confrontation baseline. This asymmetry indicates that FaLH03 undergoes extensive transcriptional reprogramming when confronting FgrI349.

These PCA-level observations were confirmed quantitatively by UpSet analysis of the corresponding differentially expressed gene sets (Figure S13). FgrI349’s response to its three confrontation partners was overwhelmingly partner-specific (Figure S13A): 742 FgrI349 genes were up-regulated only against FaveI494 (of 1,054 total), 342 only against FgrPH-1 (of 502 total), and 331 down-regulated only against FaveI494 (of 1,054 total). The largest intersections were single-set categories, while shared intersections across two or more partners were comparatively minor (163, 89, 77, 75, 58, 37, 31, and 97 genes distributed in 11 categories, data not shown). FgrI349 also regulated substantially more genes against FaveI494 (1,492 total) than against FgrPH-1 (637 total, itself a blended signal) or FaLH03 (407 total), mirroring the graded separation seen in Figure 9A. The intra-specific C-FgrPH-1 comparison, representing a necessarily blended signal from both *F. graminearum* strains, was further tested against both possible baselines (SC-FgrI349 and SC-FgrPH-1) rather than just SC-FgrI349 (Figure S13B). Most of the significant DEG calls are unique to one baseline comparison, not shared between both: the four large single-set bars 402, 302, 240, and 89, largely account for the bulk of it. This observation is consistent with genuine mixing of two distinct transcriptional programs rather than a single intermediate state.

Similarly, the two *F. avenaceum* strains’ own confrontation-*versus*-self-confrontation responses were both markedly asymmetric in magnitude and largely non-overlapping in identity (Figure S13c): FaLH03 differentially expressed roughly twice as many genes (3,180 total) as FaveI494 (1,527 total), consistent with the far larger PCA displacement observed for FaLH03. The four largest intersections were again single-strain categories (1,614 FaLH03-down-alone, 991 FaLH03-up-alone, 532 FaveI494-down-alone, and 420 FaveI494-up-alone). Together, these results indicate that the two *F. avenaceum* isolates respond to the same opponent through substantially different transcriptional programs rather than a shared, species-wide confrontation signature, which echoes, at the transcriptional level, the strain-specific divergence already documented morphologically and metabolomically.

GO enrichment of FgrI349’s transcriptional response to each confrontation partner (Figure S14) revealed a pattern that tracked opponent identity rather than confrontation status generically. In the intra-specific pairing, genes down-regulated in C-FgrPH-1 relative to SC-FgrI349 (C-FgrPH-1 *vs.* SC-FgrI349) were enriched for *secondary metabolite biosynthetic process*, while up-regulated genes were enriched for *extracellular region*, *sterol biosynthetic process* [BP], *nucleoside metabolic process*, and *kinesin complex* (Figure S14A). The same comparison evaluated against FgrPH-1’s own self-confrontation baseline (C-FgrPH-1 *vs.* SC-FgrPH-1) showed a consistent pattern : *secondary metabolite biosynthetic process* was again enriched among down-regulated genes, while up-regulated genes were enriched for *phosphopantetheine binding* and *polysaccharide metabolic process* (Figure S14B). In both inter-specific confrontations, by contrast, *secondary metabolite biosynthetic process* was enriched among up-rather than down-regulated genes. Against FaveI494, it was the most significantly enriched term overall, alongside *oxidoreductase activity*, *transmembrane transport*, *transmembrane transporter activity*, and *extracellular region* among up-regulated genes; down-regulated genes were enriched for *metabolic process* and *catalytic activity* (Figure S14C). Against FaLH03, up-regulated genes were significantly enriched exclusively for *secondary metabolite biosynthetic process* and *oxidoreductase activity*, with no significantly enriched terms detected among down-regulated genes (Figure S14D). Thus, FgrI349 down-regulates secondary metabolite biosynthesis in the intra-specific confrontation but up-regulates the same functional category in both confrontations with *F. avenaceum*.

GO enrichment of the two *F. avenaceum* strains’ own transcriptional responses to confrontation with FgrI349, relative to each strain self-confrontation baseline, revealed contrasting patterns between the two isolates (Figure S15). In FaveI494, no GO terms were significantly enriched among up-regulated genes, whereas down-regulated genes were enriched for a coherent module of cell-division-and chromosome-segregation-related categories (*e.g.*, *nuclear division*, *mitotic nuclear division*, *organelle fission*) together with *cell surface*, *extracellular region*, *fungal-type cell wall*, and *spindle microtubule* (Figure S15A). In FaLH03, up-regulated genes were enriched predominantly for core metabolic categories *(e.g., carboxylic acid metabolic process*, *oxoacid metabolic process*, and *small molecule metabolic process*) together with *oxidoreductase activity*; down-regulated genes were enriched for *plasma membrane and cell periphery*, along with several transport-and cell-wall-associated categories, including *fluconazole transmembrane transporter activity*, *fluconazole transport*, *cell wall and external encapsulating structure*, and *alcohol transmembrane transporter activity* (Figure S15B).

## Discussion

This study provides a comprehensive multi-omic analysis of strain-specific fungal competitive interactions, integrating transcriptomics, metabolomics, microscopy, and colorimetric profiling to reveal that closely related fungal pathogens deploy fundamentally different molecular strategies during interspecific encounters. Our findings challenge the traditional view that competitive responses are conserved at the species level and demonstrate that intraspecific variation can rival interspecific differences in determining competitive outcomes.

### Monoculture growth and metabolic specialization: the foundation of competitive capacity

Both *F. graminearum* strains grew substantially faster than either *F. avenaceum* strain in monoculture (Figure 1, Figure S1), and were metabolomically characterized by maltose and proline betaine (Figure 3). Proline betaine is a fungal osmoprotectant linked to active betaine methylation (Inoue et al. 2024), while maltose metabolism is a well-characterized regulatory node for growth induction in filamentous fungi and yeasts alike (Wang et al. 2002; Ichikawa et al. 2021). Together with the monoculture GO enrichment, *i.e.,* down-regulation of ribosome biogenesis and rRNA processing alongside up-regulation of secondary metabolism and transport in the faster-growing FgrPH-1 (Figure S4A), our observations suggest the growth advantage of *F. graminearum* may reflect an efficient sugar-uptake/osmotic-buffering axis rather than a simple difference in translational capacity.

*F. avenaceum* strains, by contrast, diverged from each other far more sharply than the two *F. graminearum* strains did, both metabolomically (FaLH03’s enniatin/lateropyrone signature *vs*. FaveI494’s distinct, largely unannotated profile; Figure 3) and transcriptionally (Figure 4B), despite comparable growth rates between them. This dissociation between growth rate and transcriptional/metabolic identity is consistent with adaptive metabolic specialization documented elsewhere in filamentous fungi (Brakhage 2013; Keller 2019), and is a first hint of the strain-level divergence observed throughout this study.

### Self-recognition and conspecific interaction: minimal morphological perturbation but a partner-dependent transcriptional signature

Self-confrontation (SC) provoked essentially no morphological or metabolomic reprogramming in any strain relative to monoculture (Figure 5, Figures S5, Figure S6), providing a clean baseline against which to interpret confrontation responses. Transcriptionally, however, the *F. graminearum* SC comparisons showed that much of the divergence between FgrPH-1 and FgrI349 is constitutive, present already in monoculture and largely unchanged in self-confrontation (Figure 6A), rather than induced by confrontation itself.

In the case of the intra-specific confrontation between FgrI349 and FgrPH-1, because the sample necessarily contains a blended signal from both conspecific strains analyzed against a shared reference genome, we evaluated it against each strain’s own baseline separately, which yielded different functional readouts depending on which baseline was used (Figure S14A and S14B): a modest induction of nucleoside metabolism and sterol biosynthesis relative to FgrI349’s baseline (consistent with active growth/membrane remodeling), against otherwise consistent down-regulation of secondary metabolism relative to either baseline. That the two baseline comparisons disagreed on which specific genes were involved (Figure S13B) supports genuine mixing of two distinct transcriptional programs in the sample rather than a single intermediate state. Nonetheless, although significant, this self-and intraspecific transcriptional signature remained modest in scale relative to the interspecific responses, consistent with these fungi investing comparatively little in reprogramming against genetically similar individuals while maximizing competitive responses against heterospecific competitors.

### Interspecific competition: visual, chemical, and molecular signatures of antagonism

Competitive rather than synergistic interactions are generally accepted to occur among *Fusarium* species during the infection process (Dweba et al. 2017; Petrucci et al. 2023). The molecular mechanisms underlying these competitive dynamics have remained poorly understood. Our confrontation assays revealed opponent-specific morphological and chemical signatures at the interaction zone (Figure 7, Figure S8, Figure S9). FgrI349 showed consistent metabolic/biosynthetic activation against both *F. avenaceum* strains, while the two *F. avenaceum* strains themselves diverged from each other, FaLH03 shifting in parallel with FgrI349, FaveI494 shifting in the opposite direction, and FgrI349’s own hue response differing depending on which partner it faced. This colorimetric divergence between FaLH03 and FaveI494 is the first cue of the large strain-level split observed in this study.

The metabolomic data anchor these visual signals to specific chemistry: enniatin B/B1 and lateropyrone (putative) remained strongly and specifically associated with FaLH03 across confrontation contexts (Figure 8D). Enniatin B, a toxic cyclohexadepsipeptide secondary metabolite (Jestoi 2008; Urbaniak et al. 2020), has been reported to inhibit *F. graminearum* while promoting *F. avenaceum* growth (Beccari et al. 2017; Ederli et al. 2021a), raising the possibility that these compounds function less as plant-virulence factors and more as direct competitive weapons against co-occurring *Fusarium* species. *F. graminearum*, in contrast, maintained a comparatively stable metabolomic identity (maltose, proline betaine) across confrontation scenarios, suggesting a more conservative chemical strategy than the metabolically dynamic *F. avenaceum* strains.

This chemical asymmetry has a growth-level correlate: FgrI349’s own growth increased specifically in interspecific confrontations but not against its conspecific FgrPH-1 (Figure 8A). This observation is consistent with reports that *F. graminearum* regulates killer-toxin genes during competitive interactions with other *Fusarium* species (Petrucci et al. 2025), and suggests a recognition mechanism that distinguishes conspecific from heterospecific competitors.

### Directionality of transcriptional responses: induction in *F. graminearum*, repression in *F. avenaceum*

Across every confrontation comparison, FgrI349 up-regulated substantially more genes than it down-regulated relative to its own baseline, while both *F. avenaceum* strains showed the opposite direction, down-regulated genes modestly outnumbering up-regulated ones (Figures S14 and S15). A modest 1.2–1.4-fold asymmetry of the transcriptional response can be observed for *F. avenaceum* strains, that can be interpreted as a directional tendency rather than a wholesale shutdown. Taken together, these observations point to a species-level difference in how competitive transcriptional programs are built: FgrI349’s response looks like a targeted induction layered onto a largely stable background, while the *F. avenaceum* strains’ responses lean toward modest net resource re-allocation even as specific functions are concurrently up-regulated.

A stricking example of this asymmetry lays in the significant enrichment of the GO category *secondary metabolite biosynthetic process* in every FgrI349 confrontation comparison, but with direction flipped with opponent identity: down-regulated intraspecifically, up-regulated against both *F. avenaceum* strains (Figure S14). This opponent-dependent reversal of a single biologically coherent category appears as a striking signature of competitor discrimination. Notably, this same category also showed a nominal, sub-threshold up-regulation signal on FaveI494’s own side of its confrontation with FgrI349 (see below), hinting that this may not be a purely one-sided discrimination response by FgrI349 alone.

### Strain-level variation rivals species-level differences

One striking finding of this study is that FaLH03 and FaveI494, genetically close strains of the same species facing the identical opponent, display substantially different competitive transcriptional programs. This observation is extends prior evidence of intraspecific variation shaping virulence and host range in other fungal pathogens (Parker et al. 2023) into the domain of direct fungal-fungal competition.

FaLH03 showed a focused response on core oxidative, carboxylic, and amino-acid metabolism, with reduced investment in membrane and transport machinery (Figure S15B). However, it lacked a distinct secondary metabolite biosynthesis GO signature, despite the observed enniatin/lateropyrone (putative) production at the metabolomic level.

FaveI494’s response differed in kind, not just scale: no GO terms were significant among its upregulated genes, while its downregulated genes formed a coherent cell-division/chromosome-segregation module (Figure S15A), suggesting a growth-arrest signature rather than an active enzymatic response. Nonetheless, examining the underlying statistics for this comparison revealed that secondary metabolite biosynthetic process (GO:0044550; 6 /40 genes [present/background], raw p = 7.7×10⁻⁵) and its parent term (GO:0019748; 7/49 genes [present/background], raw p = 2.6×10⁻⁵) showed nominal enrichment that did not survive multiple-testing correction (adjusted p = 0.33 and 0.22). This raises the possibility that FaveI494’s visible carmine-red pigmentation (Figure 7B) is accompanied by a modest, sub-threshold transcriptional signal on its own side of the interaction.

A striking pattern to emerge from these confrontations is a decoupling between the observation that FaveI494 elicited the largest and most strongly up-regulated response from FgrI349 of any partner (Figure S14C) and the largest displacement in principal component space (Figure 9A), and a more muted, largely repressive transcriptional footprint relative to its own self-confrontation baseline. FaLH03, conversely, showed the larger shift in its own transcriptome relative to its own baseline (Figure 9B) while eliciting the smallest response of any partner from FgrI349 (Figure S14D). An UpSet comparison of the two strains’ own response gene sets confirmed they are also largely non-overlapping at the individual-gene level (Figure S13C). Together, these results indicate that “*how much a strain itself reprograms transcriptionally*” and “*how much competitive pressure it exerts on its opponent*” are not the same axis, an important distinction for how competitive strength should be assessed in future studies.

Among hypotheses that could drive such divergent strategies between two strains of the same species such as FaveI494 and FaLH03, mobile genetic elements such as Starships, known to generate strain heterogeneity by mobilizing virulence-related genes and biosynthetic gene clusters (Gluck-Thaler et al. 2022), are possible contributors. Mycovirus infection is another hypothesis, mycoviruses being known to alter fungal growth, virulence, and metabolite production in virus-and host-strain-dependent ways (Yu, and Kim 2020; Kuroki et al. 2023). Mycovirus are common in Fusarium head-blight-associated species (Buivydaitė et al. 2024), and can measurably shift host transcription, including down-regulation of metabolic and transport genes such as those reported here, even when infection produces no obvious external phenotype (Cho et al. 2012a; Lee, Cho, Yu, Son, Choi, Min, Lee, Kim, et al. 2014).

### Opponent-specific response plasticity: tailored competitive strategies

FgrI349’s response was not a unique confrontation program scaled up or down by opponent; it differed in both magnitude and functional content depending on who was faced. The largest and most functionally elaborate response (secondary metabolism, oxidoreductase activity, membrane transport) occurred against FaveI494; the response to FaLH03 was smaller and functionally narrower. An UpSet comparison confirmed this specificity extends to the gene level, with little overlap between opponent-specific gene sets (Figure S13A and Figure S14). This partner-specific magnitude hierarchy was independently recapitulated in PCA space (Figure 9A), in colorimetric shifts (Figure S9), and in the pigmentation phenotype itself (Figure 7B). This convergence across four independent data types gives particular confidence FaveI494 represents a distinctly potent competitive challenge for FgrI349, met with a correspondingly tailored response.

### Ecological and evolutionary implications

*Fusarium* interactions during germination are known to be predominantly competitive (Wagacha et al. 2012). Our results extend this knowledge to later developmental stages and show that competitive interactions can actively amplify, rather than merely reflect, pre-existing fitness differences, as demonstrated by FgrI349’s opponent-specific growth induction. Moreover, FaveI494 provokes the strongest opponent response while exhibiting the most modest transcriptional footprint relative to its self-confrontation culture. This observation suggests that competitive strength must be evaluated from both sides of an interaction, not just one partner’s transcriptome. Such a dual perspective may better explain strain dominance patterns observed in natural or agricultural fungal communities.

*F. graminearum* documented rise to dominance in some wheat-growing regions (Nielsen et al. 2011) is consistent with the idea that such shifts can be driven by strain-level, not just species-level or environmental factors. This consideration has direct implications for biocontrol strategy. Given the demonstrated divergence between FaLH03 and FaveI494 alone, biocontrol approaches that rely on competition or antibiosis (Saravanakumar et al. 2016; Petrucci et al. 2023b) will likely need to account for intraspecific diversity in both target pathogens and biocontrol agents, rather than treating either species as a uniform competitive entity.

### Comparative context and future directions

These results argue that fungal competitive interactions are best understood as strain-specific rather than species-uniform, and that morphological, chemical, and molecular responses, while coordinated, are not always in lockstep. Several extensions follow directly from this work. Mechanistically, the rapid, coordinated transcriptional shifts we observe are consistent with chromatin-level regulation (Pfannenstiel, and Keller 2019), and testing for epigenetic or small-RNA involvement would help distinguish transcription-factor-driven from chromatin-driven competitive responses. Ecologically, our pairwise design cannot capture the frequency-dependent dynamics or emergent hierarchies likely to arise in multispecies communities, which will require significantly more complex confrontation designs. Temporally, our data are single-timepoint snapshots of an inherently dynamic process. Time-series sampling, including earlier stages before hyphal contact, could resolve the sequence in which morphological, chemical, and transcriptional responses actually unfold (though such sampling in time and space is not trivial).

## Conclusion

This study demonstrates that fungal competitive interactions involve sophisticated, strain-specific molecular programs that challenge simple species-level generalizations. FgrI349 consistently mounted a predominantly inductive transcriptional response, graded in magnitude and distinct in functional content by opponent, with secondary metabolite biosynthesis itself flipping regulatory direction between intraspecific and interspecific encounters. The two *F. avenaceum* strains, by contrast, showed modest net down-regulation in their own responses but pursued markedly different, largely non-overlapping strategies: FaLH03 shifted more in its own transcriptome, centered on core oxidative metabolism, while FaveI494 showed a more contained, cell-cycle-centered response yet provoked by far the strongest reaction from its opponent. This decoupling between a strain’s own transcriptional reprogramming and the competitive pressure it exerts on its opponent likely is a generalizable observation for how competitive strength should be assessed in future studies of this kind.

The integration of morphological, colorimetric, metabolomic, and transcriptomic data shows that visual, chemical, and molecular signals of fungal competition are coordinated, but not always strictly correlated across data types. Future work integrating genomics, epigenomics, temporal dynamics, and community ecology will be needed to fully understand the evolution and ecology of fungal competitive interactions.

## Acknowledgement

The authors thank Vance C. Huskins for his careful proofreading. The work showed here was supported jointly by the department the INRAE departments SPE (Plant Health and Environment) and MICA (Microbiology of Food Chain). Mobility fellowships to perform all microscopy work in the lab of N. Mach were awarded to M. Navarro by the University of Bordeaux and the INRAE network REacTION.

## Data availability

RNA-seq reads were deposited at the European Nucleotide Archive under the project accession number PRJEB88188. Metabolomics data can be retrieved from the public repository Recherche Data Gouv (https://doi.org/10.57745/AB4UN6).

## Supplementary Figures & figure legends

**Supplementary Figure S1.**
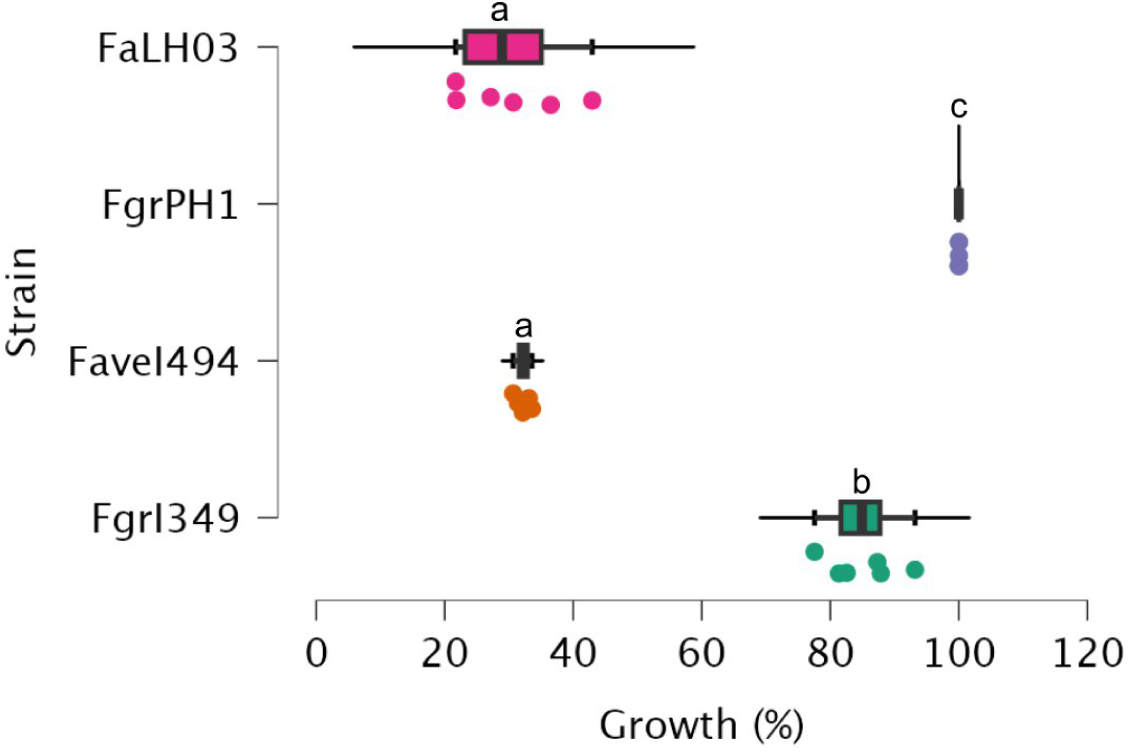
Fungal growth measured as the surface of the Petri dish occupied by mycelium, in percentage of the total area. The raincloud plot displays the distribution (n = 6 per strain) of the percentage growth measured for each of the four fungal strains. The x-axis represents the growth percentage, and the y-axis lists the strains. The letters a,b, and c indicate the statistical groupings based on Kruskal-Wallis test (non-parametric ANOVA) followed by Dunn’s Post Hoc pairwise comparisons and Benjamini-Hochberg correction (p < 0.05).

**Supplementary Figure S2.**
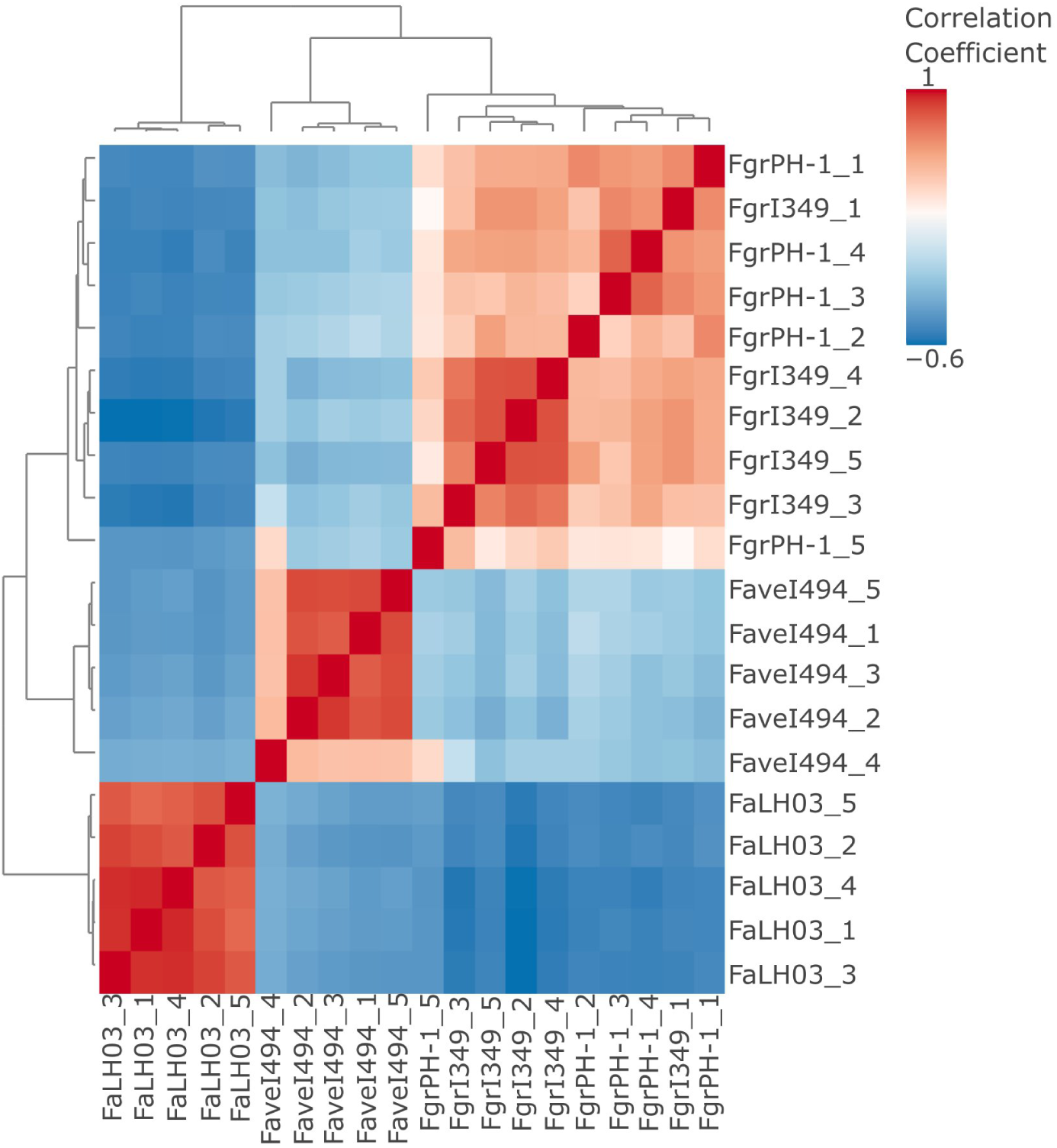
Correlation (Pearson) matrix between different samples metabolomic profiles. The color scale, ranging from -0.6 to 1 represents the strength and direction of the correlations. The dendrograms on the axes indicate hierarchical clustering of the samples, grouping those with similar correlation patterns.

**Supplementary Figure S3.**
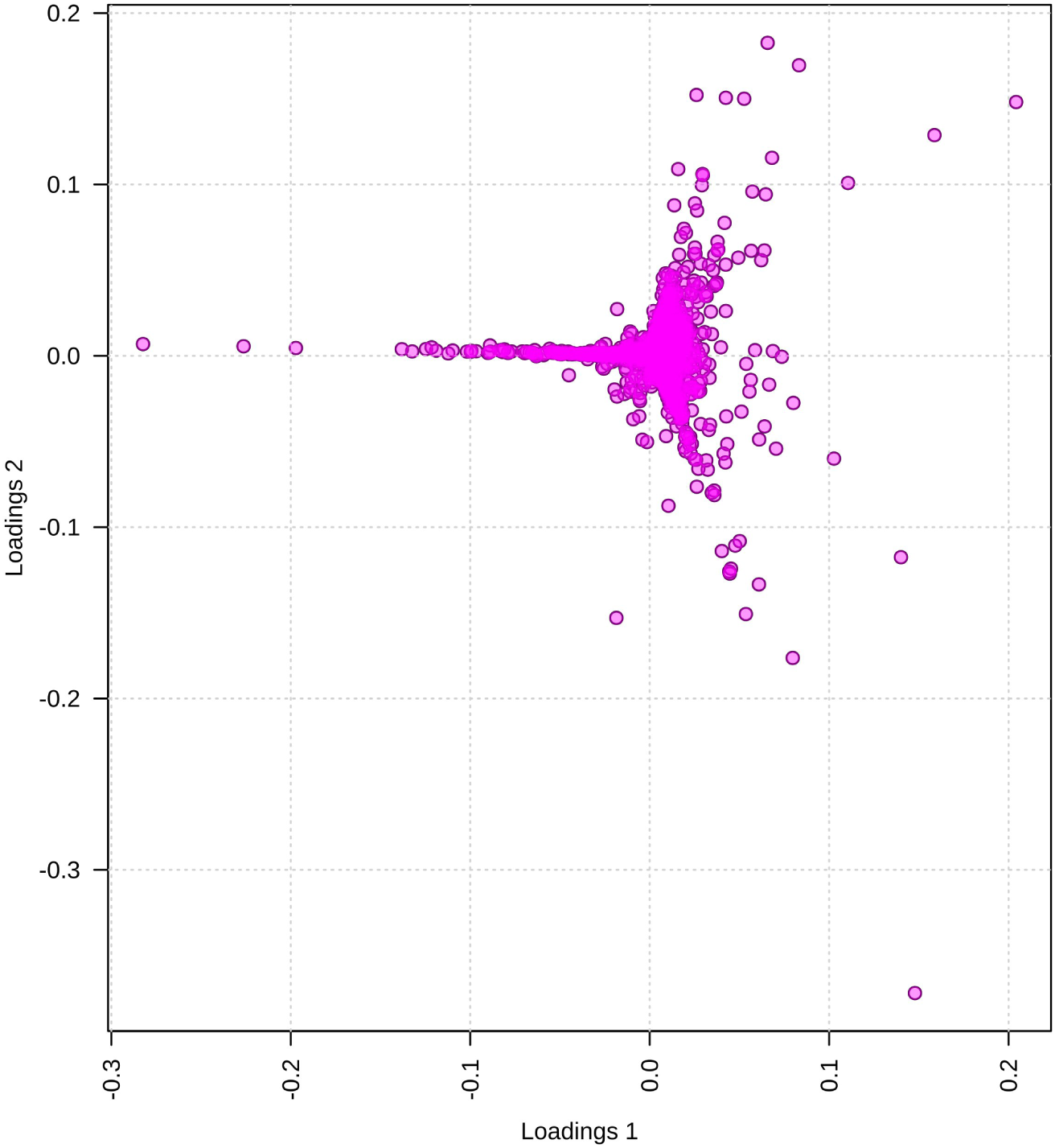
PCA loadings plot of metabolomic profiles.

**Supplementary Figure S4.**
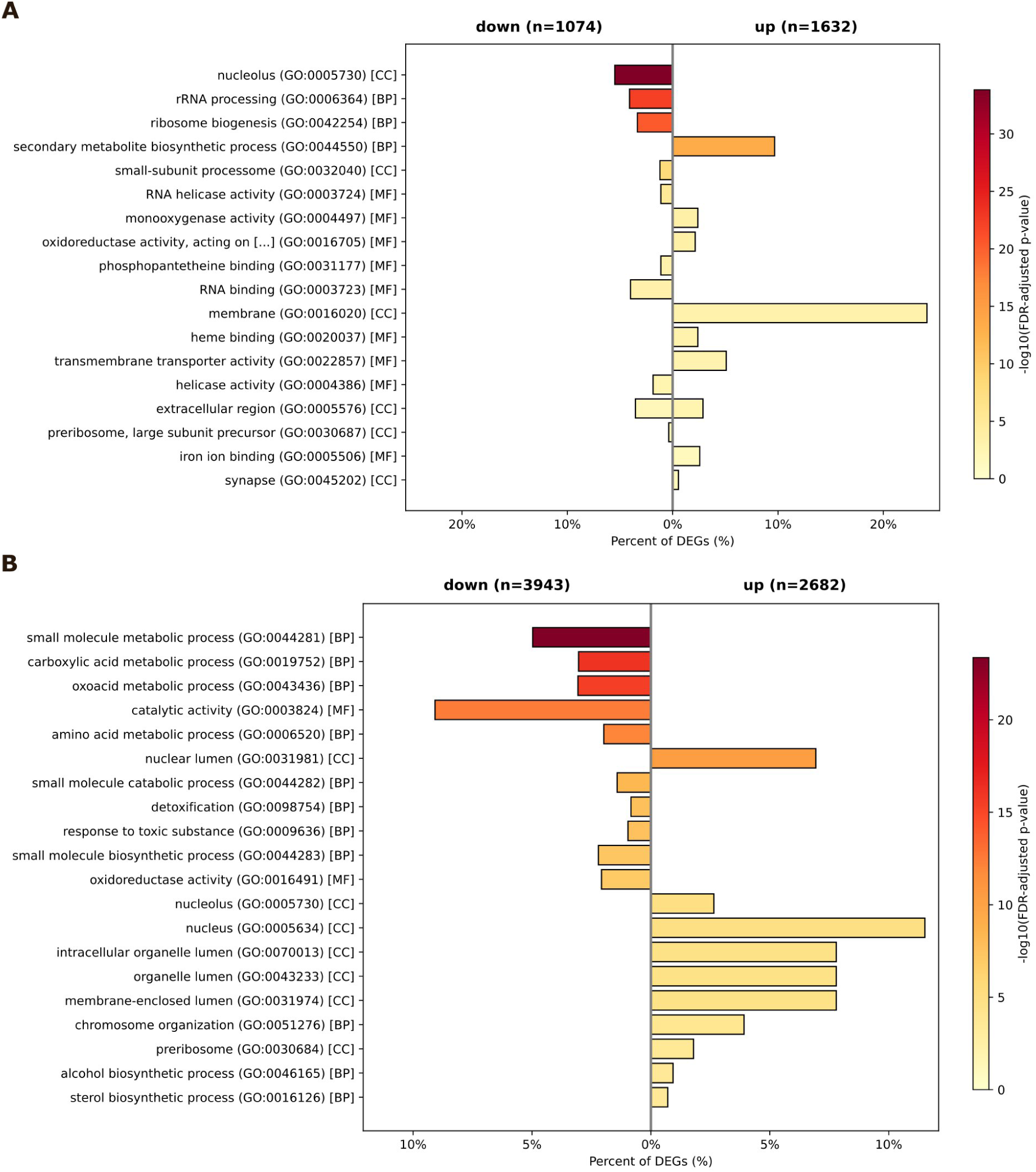
Gene Ontology (GO) term enrichment analysis of differentially expressed genes *F. graminearum* FgrPH-1 and FgrI349 **(A)**, and *F. avenaceum* FaveI494 and FaLH03 **(B)**. Horizontal bar plot showing enriched GO categories for upregulated (right side) and downregulated (leftside) genes. The length of each bar indicates the percentage of genes annotated to the respective GO term, while color intensity represents the statistical significance of enrichment expressed as -log₁₀(adjusted p-value).

**Supplementary Figure S5.**
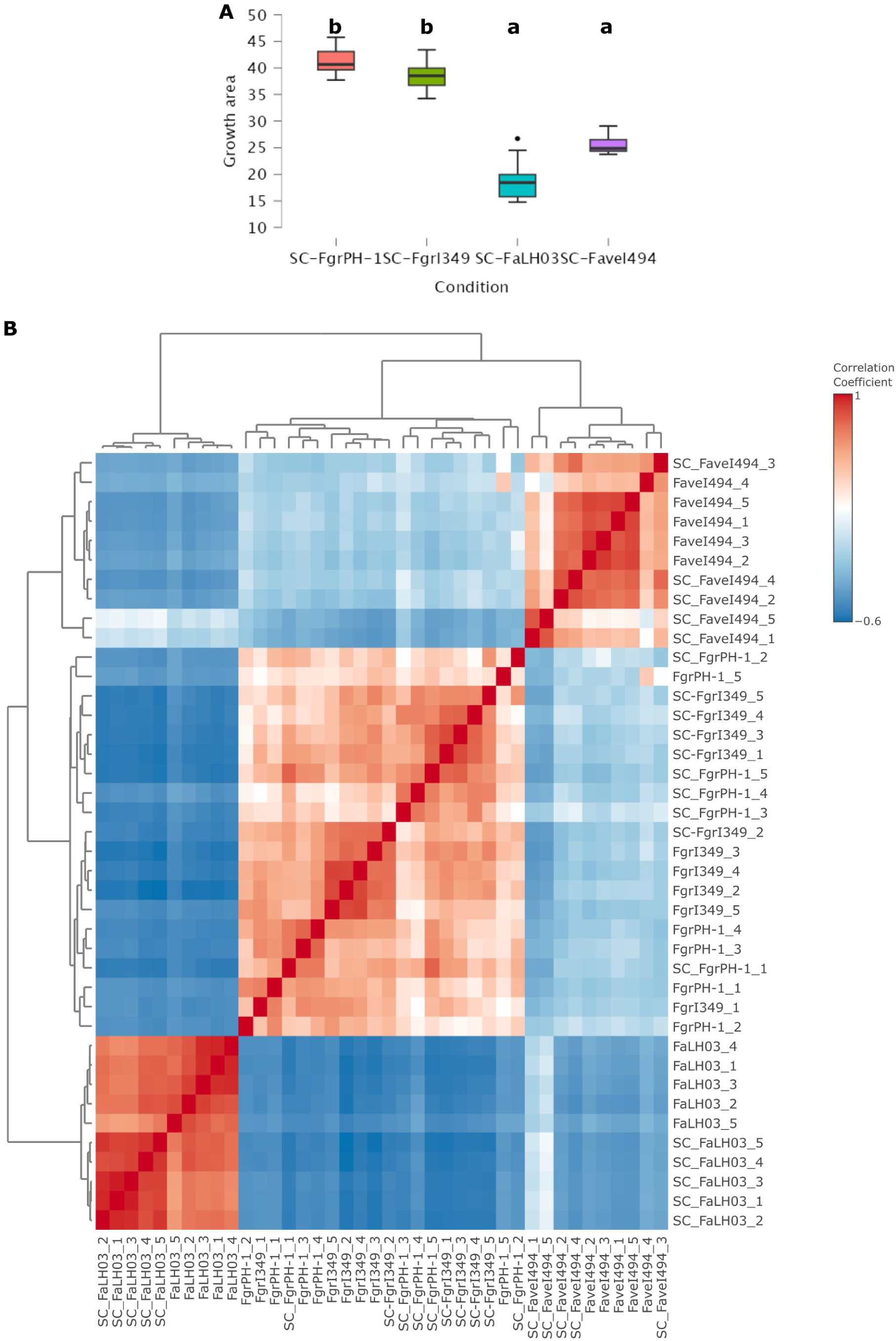
Characteristics of self-confrontation cultures. **(A)** Fungal growth measured as the surface of the Petri dish occupied by mycelium for each interacting partner, in percentage of the total area (n = 12 per strain, *i.e.,* six confrontations of two identical partners per Petri dish). The letters a and b indicate the statistical groupings based on Kruskal-Wallis test (non-parametric ANOVA) followed by Dunn’s Post Hoc pairwise comparisons and Benjamini-Hochberg correction (p < 0.05). **(B)** Correlation (Pearson) matrix between different SC and monoculture samples metabolomic profiles. The color scale, ranging from -0.6 to 1 represents the strength and direction of the correlations. The dendrograms on the axes indicate hierarchical clustering of the samples, grouping those with similar correlation patterns.

**Supplementary Figure S6.**
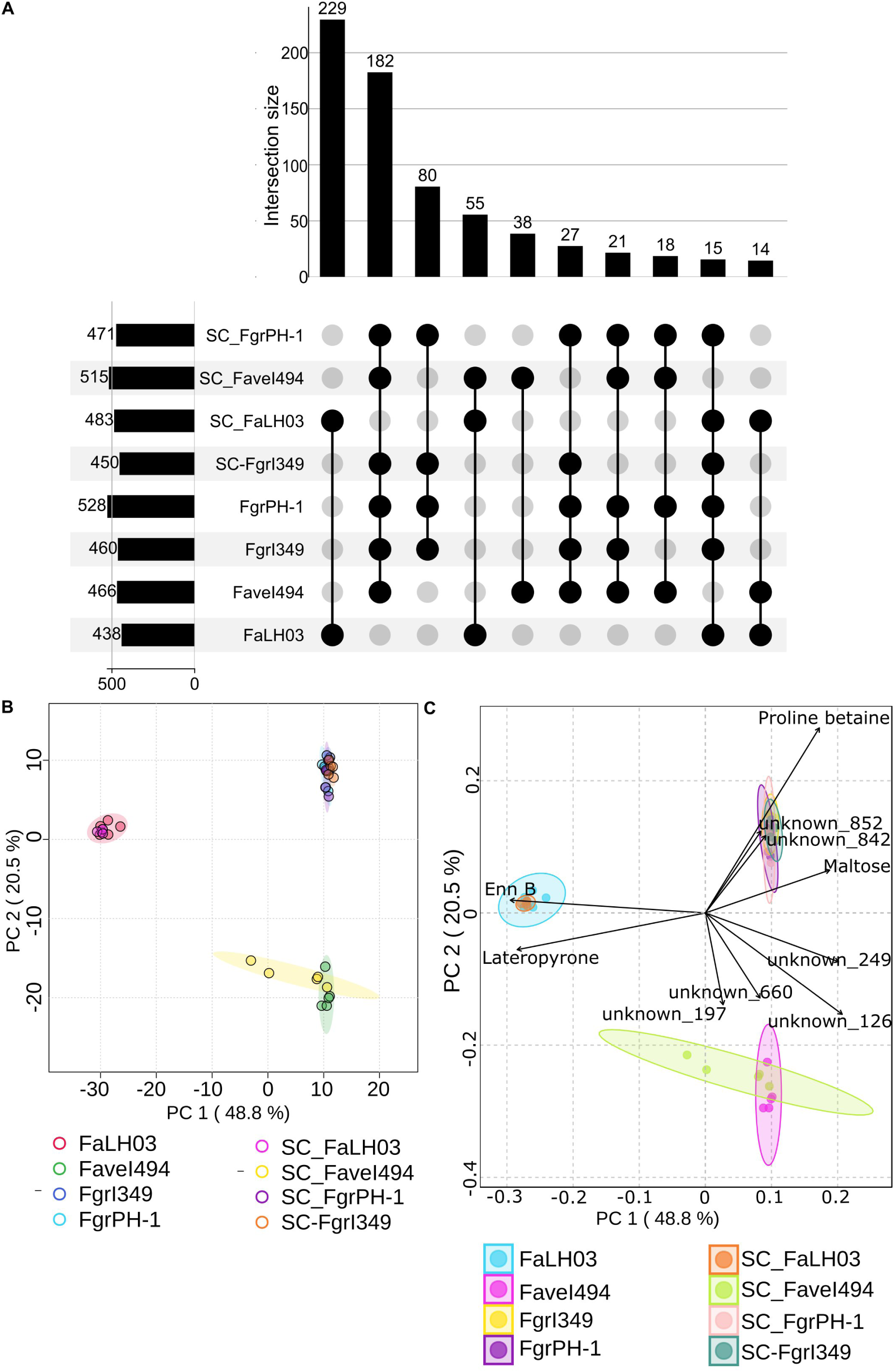
Metabolomic profiling of strains grown in monocultures and selfconfrontation (SC) **(A)** Upset Plot visualizing the Intersection of metabolomic features across strains and conditions. The bar plot (top) quantifies the number of unique and shared metabolomic features among the samples, with only the top 10 intersections displayed for clarity. The connected dots (bottom) indicate the specific intersections between the sets. **(B)** PCA scores plot of metabolomic profiles of *F. avenaceum* FaLH03 and FaveI494, and *F. graminearum* FgrI349 and FgrPH-1, grown in monocultures and self-confrontation (SC) conditions. The ellipses around each group represent the 95% confidence interval for the distribution of samples within that group. **(C)** PCA biplot showing the top 10 (magnitude) loading vectors. SC_FgrPH-1: self-confrontation of *F. graminearum* FgrPH-1; SC_FaveI494: self-confrontation of *F. avenaceum* FaveI494.; SC_FaLH03: self-confrontation of *F. avenaceum* FaLH03; SC_FgrI349: self-confrontation of *F. graminearum* FgrI349; FgrPH-1, FgrI349, FaveI494, FaLH03: Monocultures of the respective strains.

**Supplementary Figure S7.**
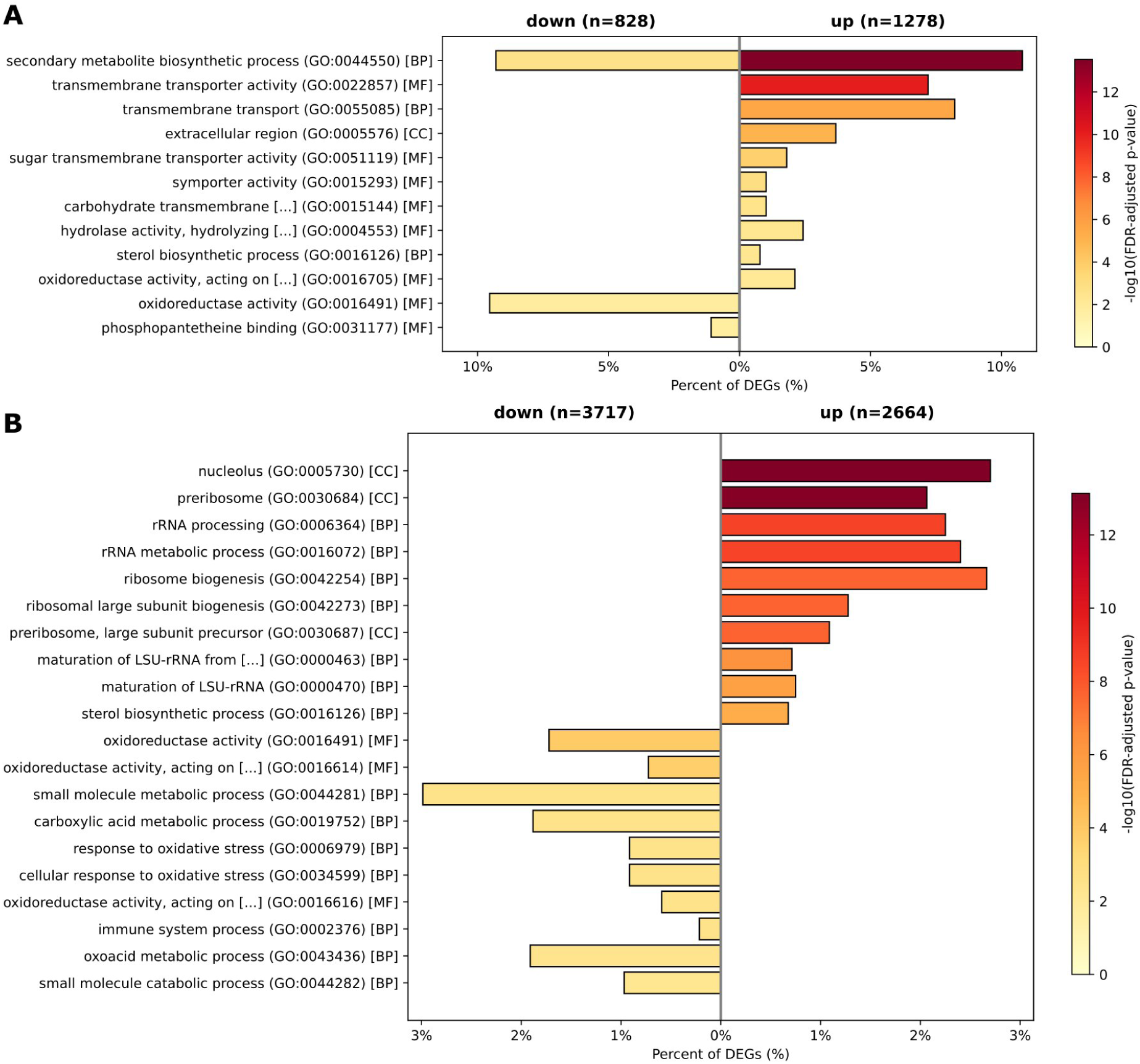
Gene Ontology (GO) term enrichment analysis of differentially expressed genes between *F. graminearum* strains FgrPH-1 and FgrI349 **(A)**, and between *F. avenaceum* strains FaveI494 and FaLH03 **(B)**, under self-confrontation. Horizontal bar plots show enriched GO categories for up-regulated (right side) and down-regulated (left side) genes. The length of each bar indicates the percentage of differentially expressed genes annotated to the respective GO term, while color intensity represents the statistical significance of enrichment expressed as -Log₁₀(adjusted p-value).

**Supplementary Figure S8.**
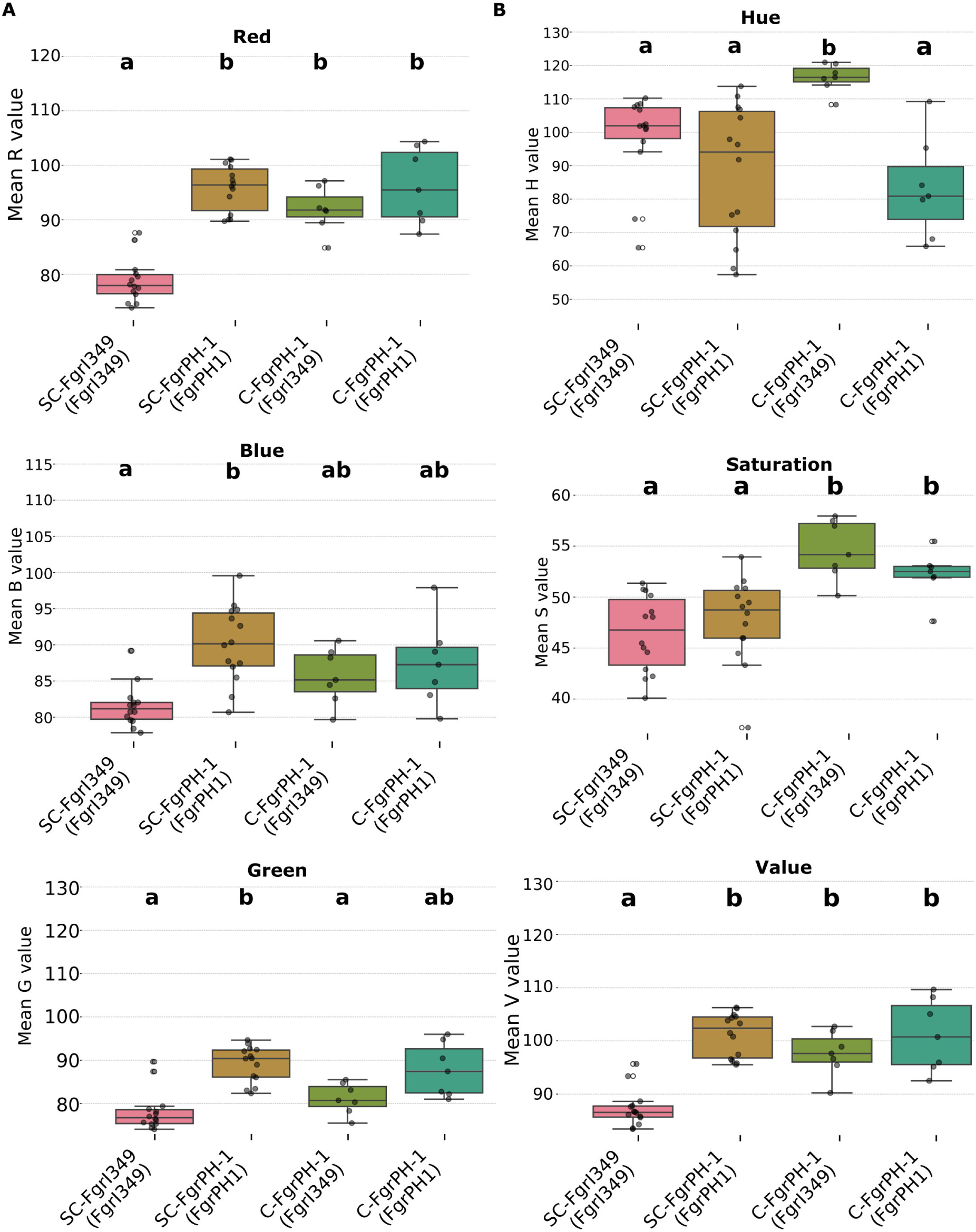
Comparison of RGB and HSV distributions across *F. graminearum* FgrI349 and FgPH-1, in confrontation with FgrI349 or self-confrontations. **(A)** Distribution of RGB color values (Red, Green, Blue) and **(B)** HSV color parameters (Hue reflecting color type, Saturation reflecting color intensity, Value reflecting brightness). C-FgrPH-1: FgrI349 *vs.* FgrPH-1; the strain name in parenthesis is the considered interacting partner. The letters a,b,c, and d indicate the statistical groupings based on Kruskal-Wallis test (non-parametric ANOVA) on per-image aggregated mean pixel values (n=12) followed by Mann-Whitney U pairwise comparison and multiple comparison Benjamini-Hochberg correction results (p < 0.05). Whiskers extend to 1.5× the interquartile range, with outliers shown as individual points.

**Supplementary Figure S9.**
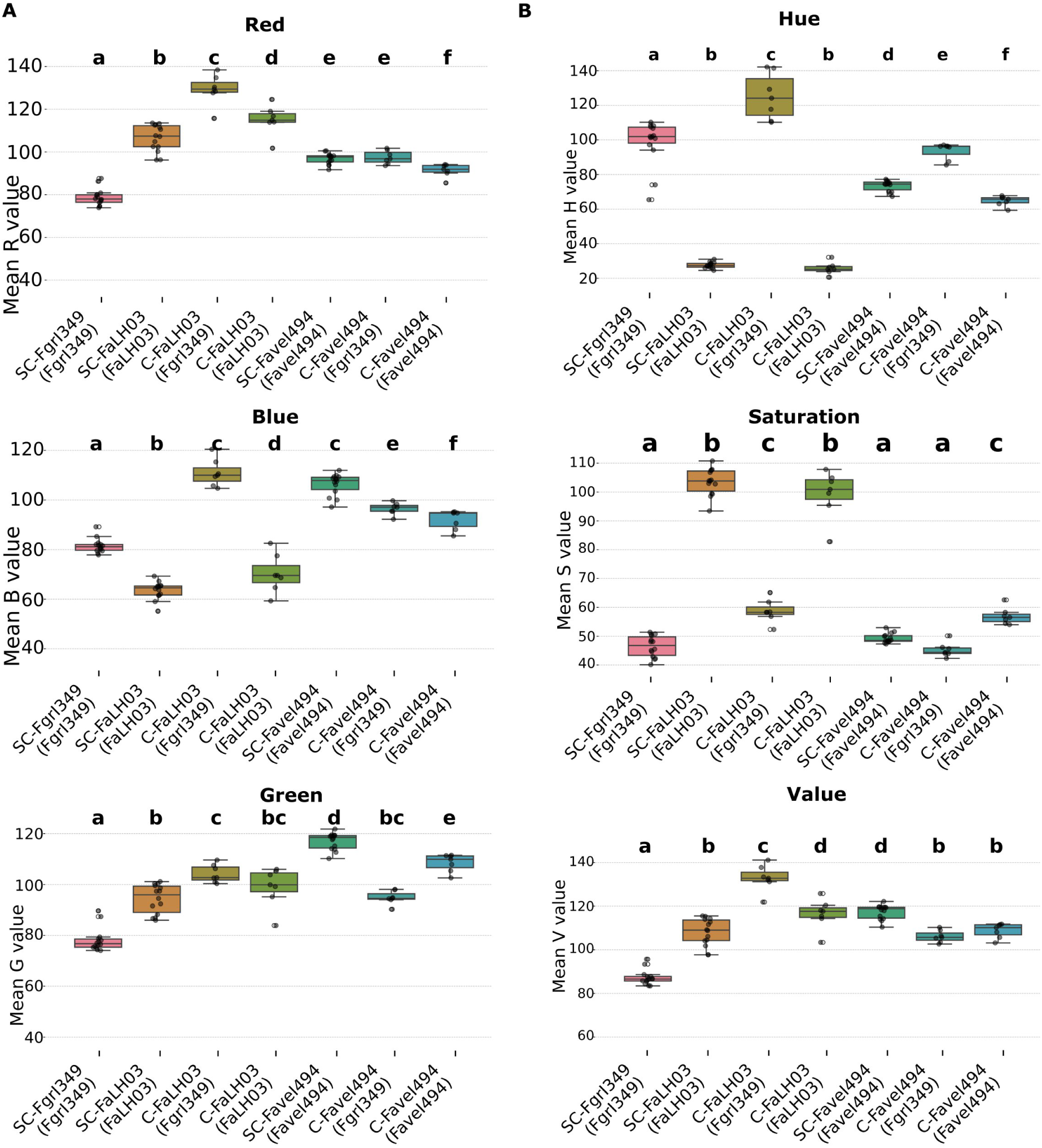
Comparison of RGB and HSV distributions across *F. avenaceum* FaveI494 and FaLH03, in confrontation with FgrI349 or self-confrontations. **(A)** Distribution of RGB color values (Red, Green, Blue) and **(B)** HSV color parameters (Hue reflecting color type, Saturation reflecting color intensity, Value reflecting brightness). C-FaLH03 FgrI349 *vs.* FaLH03; C-FaveI494 FgrI349 *vs.* FaveI494; the strain name in parenthesis is the considered interacting partner. The letters a,b,c, and d indicate the statistical groupings based on Kruskal-Wallis test (non-parametric ANOVA) on per-image aggregated mean pixel values (n=12) followed by Mann-Whitney U pairwise comparison and multiple comparison Benjamini-Hochberg correction results (p < 0.05). Whiskers extend to 1.5× the interquartile range, with outliers shown as individual points.

**Supplementary Figure S10.**
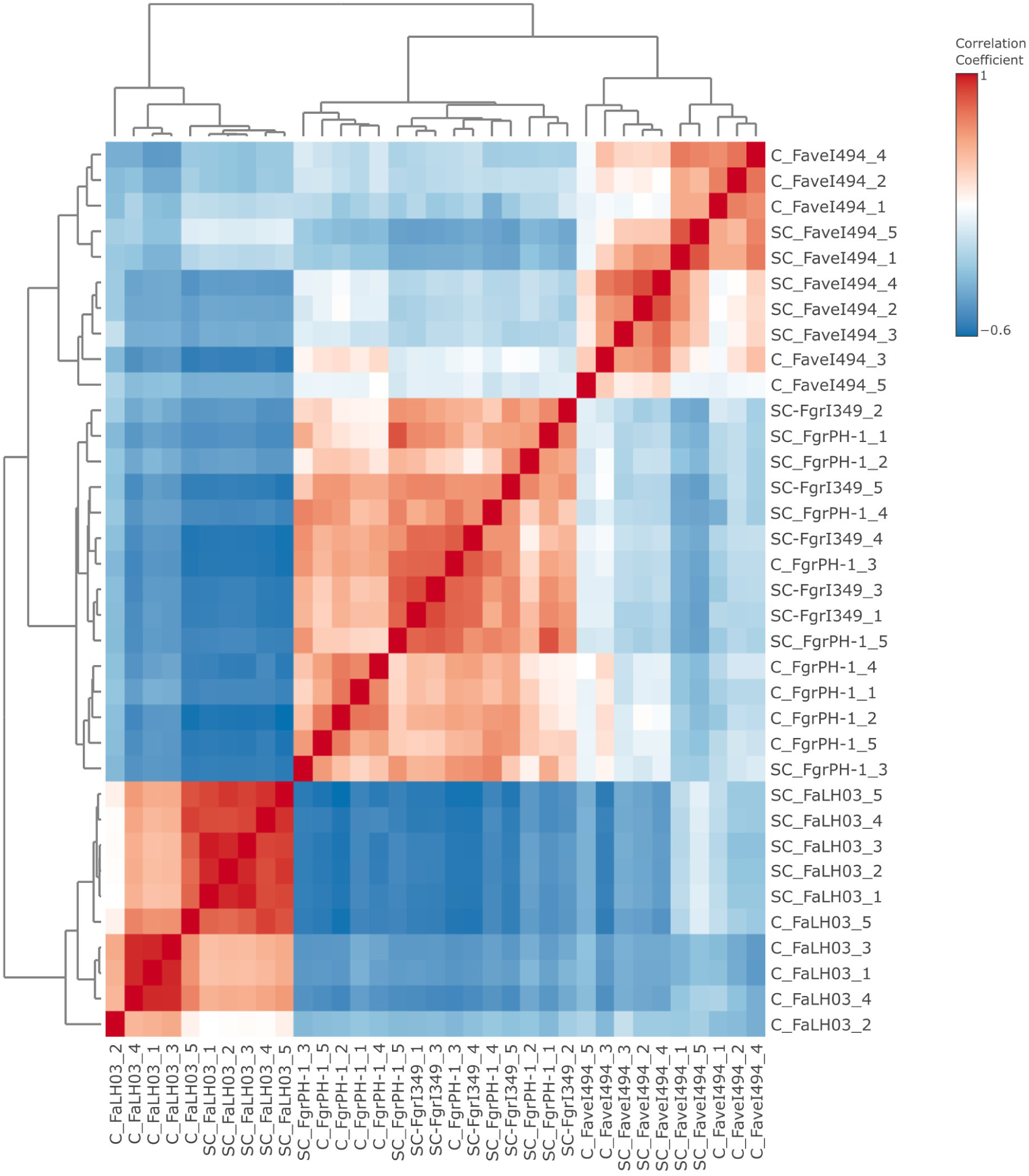
Correlation (Pearson) matrix between different SC and C samples metabolomic profiles. The color scale, ranging from -0.6 to 1 represents the strength and direction of the correlations. The dendrograms on the axes indicate hierarchical clustering of the samples, grouping those with similar correlation patterns.

**Supplementary Figure S11.**
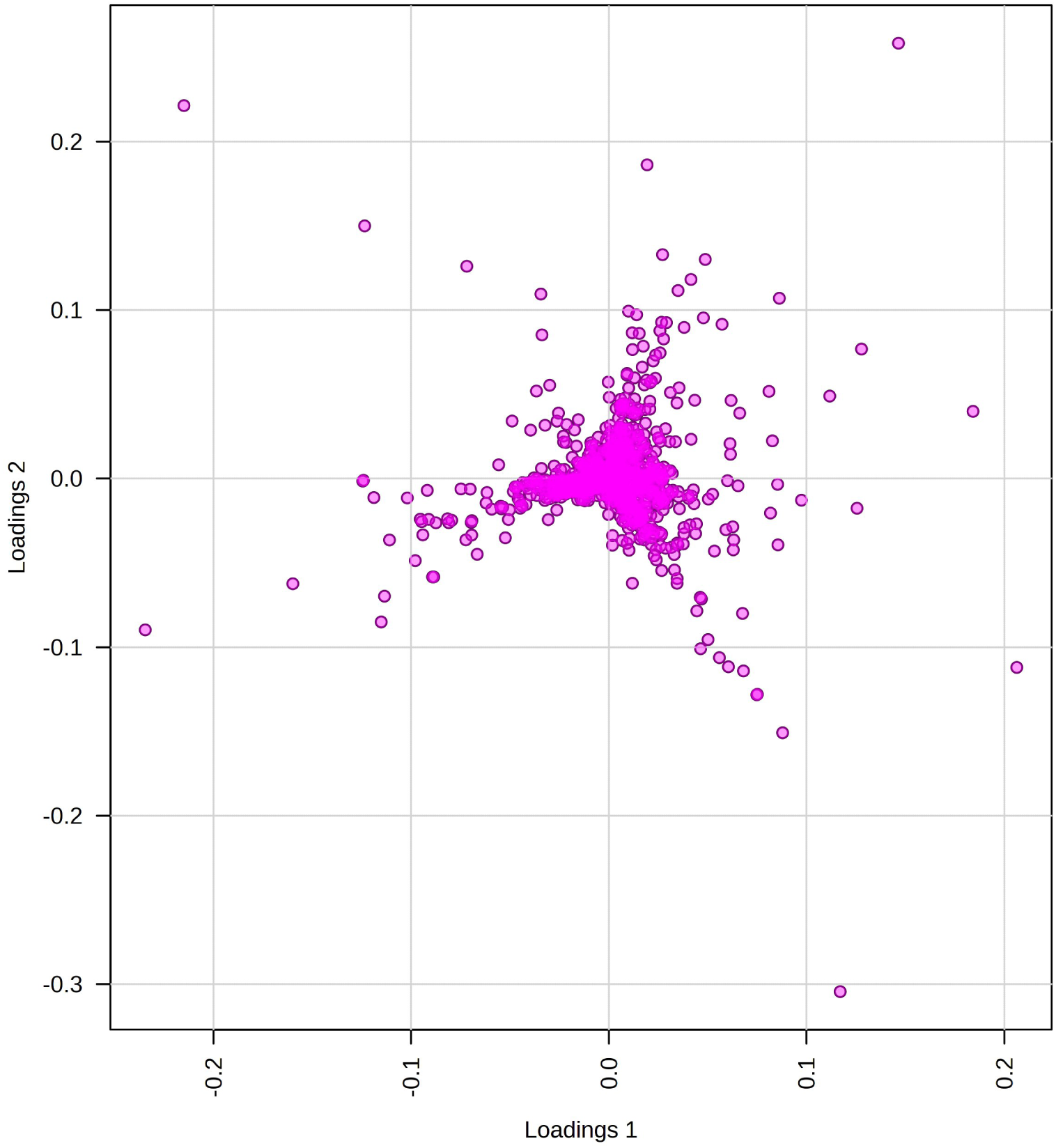
PCA loadings plot of metabolomic profiles.

**Supplementary Figure S12.**
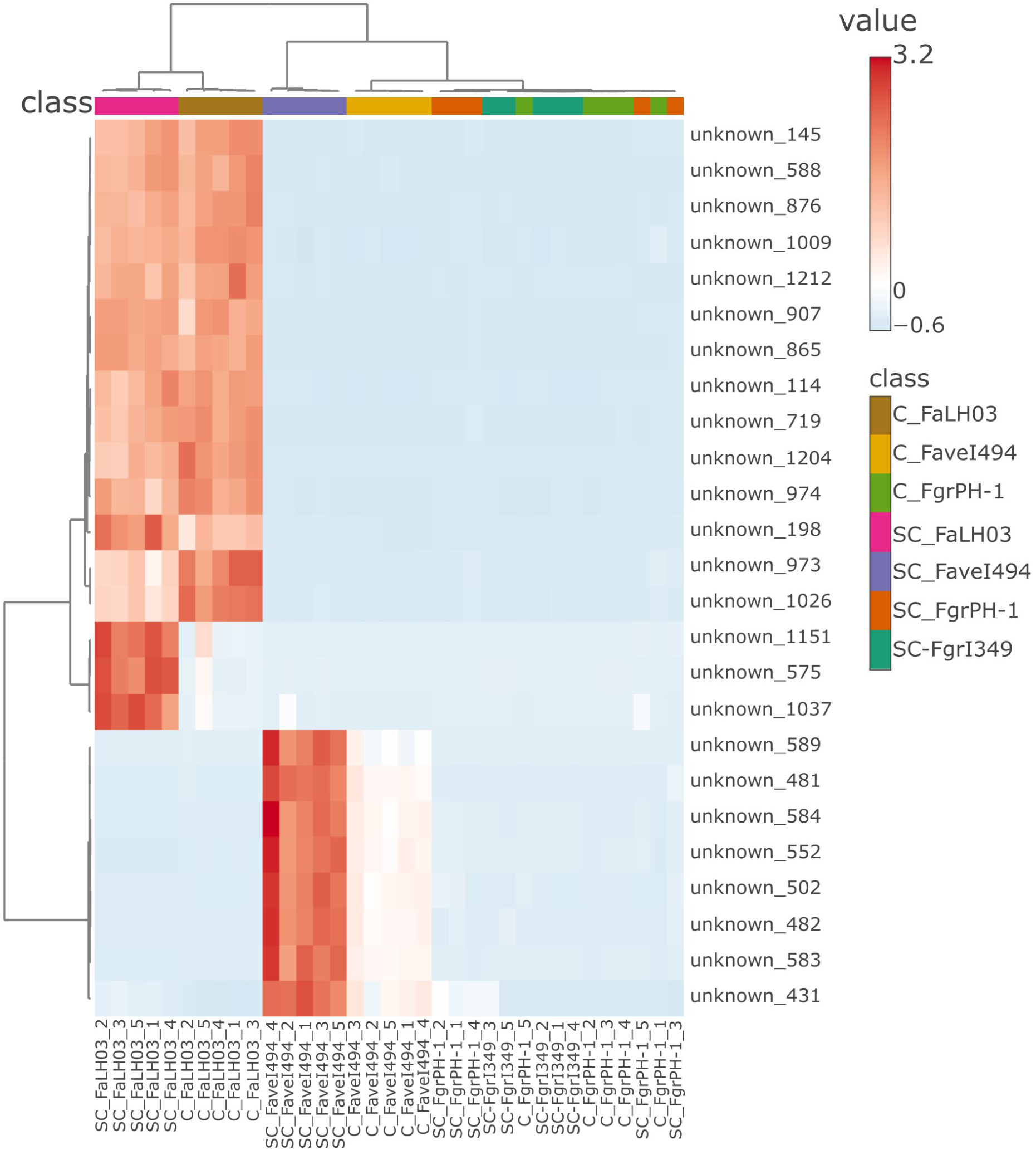
Hierarchical clustering heatmap showing the relative abundance of metabolic features across the seven confrontation or self-confrontation samples: C_FaLH03 (brown), C_Favel494 (yellow), C_FgrPH-1 (green), SC_FaLH03 (pink), SC_Favel494 (purple), SC_FgrPH-1 (red), and SC_FgrI349 (teal). Rows represent individual metabolic features, and columns correspond to biological replicates within each sample group. The color scale indicates normalized values (Z-scores), where red denotes higher relative abundance and blue denotes lower values. Both rows and columns were clustered using Euclidean distance and ward linkage. The dendrogram at the top shows sample clustering based on similarity, while the dendrogram on the left groups features that share similar abundance profiles. Only the 50 most variable features are shown. The color bar on the right represents autoscaled values from approximately −1.6 (lower abundance) to +3.2 (higher abundance).

**Supplementary Figure S13.**
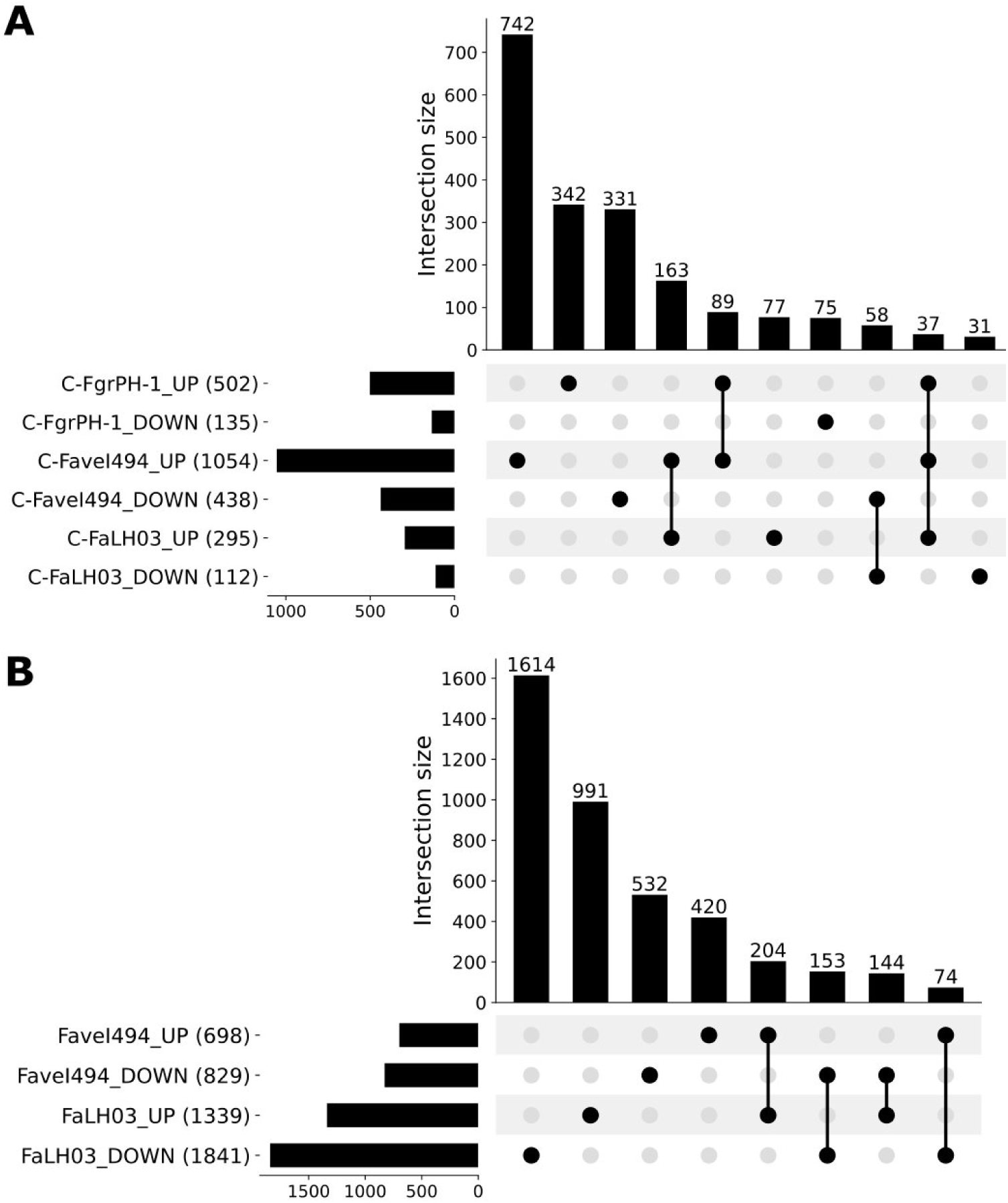
UpSet plots of differentially expressed gene sets across confrontation comparisons. **(A)** Overlap of genes differentially expressed in FgrI349’s transcriptional response to each of its three confrontation partners (FgrPH-1, FaveI494, and FaLH03), relative to FgrI349’s self-confrontation baseline, shown separately for up-and down-regulated genes. **(B)** Overlap of genes differentially expressed in the intra-specific confrontation (C-FgrPH-1), evaluated against both possible self-confrontation baselines (SC-FgrI349 and SC-FgrPH-1), shown separately for up-and down-regulated genes. **(C)** Overlap of genes differentially expressed in FaveI494’s and FaLH03’s own transcriptional responses to confrontation with FgrI349, relative to each strain’s self-confrontation baseline, shown separately for up-and down-regulated genes. Horizontal bars indicate the total number of genes in each set; vertical bars and connected dots indicate the number of genes in each intersection.

**Supplementary Figure S14.**
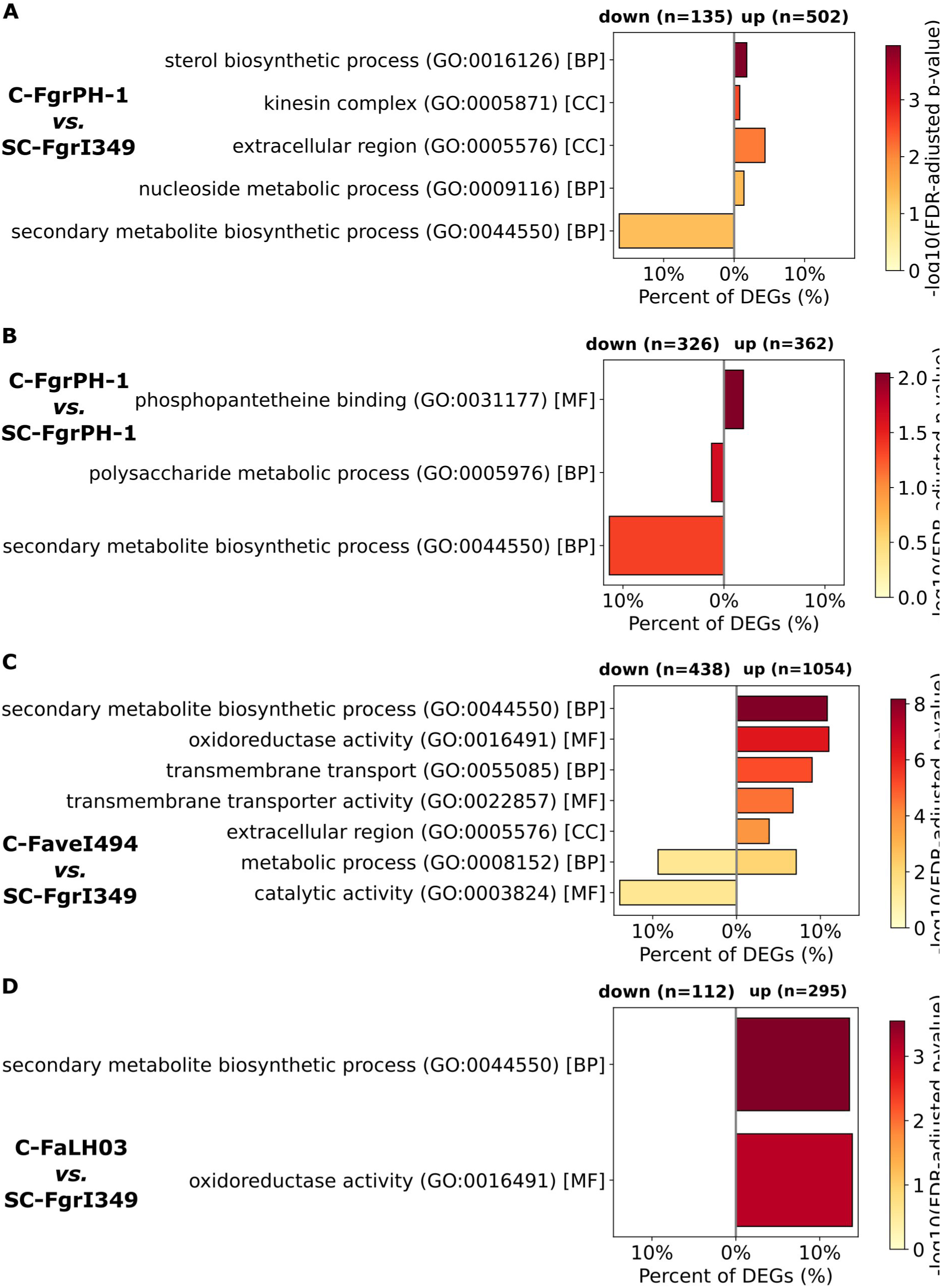
Gene Ontology (GO) term enrichment analysis of FgrI349’s differentially expressed genes across confrontation partners. **(A)** C-FgrPH-1 *vs*. SC-FgrI349. **(B)** C-FgrPH-1 *vs*. SC-FgrPH-1. **(C)** C-FaveI494 *vs*. SC-FgrI349. **(D)** C-FaLH03 *vs*. SC-FgrI349. Horizontal bar plots show enriched GO categories for up-regulated (right side) and down-regulated (left side) genes. The length of each bar indicates the percentage of differentially expressed genes annotated to the respective GO term, while color intensity represents the statistical significance of enrichment expressed as -Log₁₀(adjusted p-value).

**Supplementary Figure S15.**
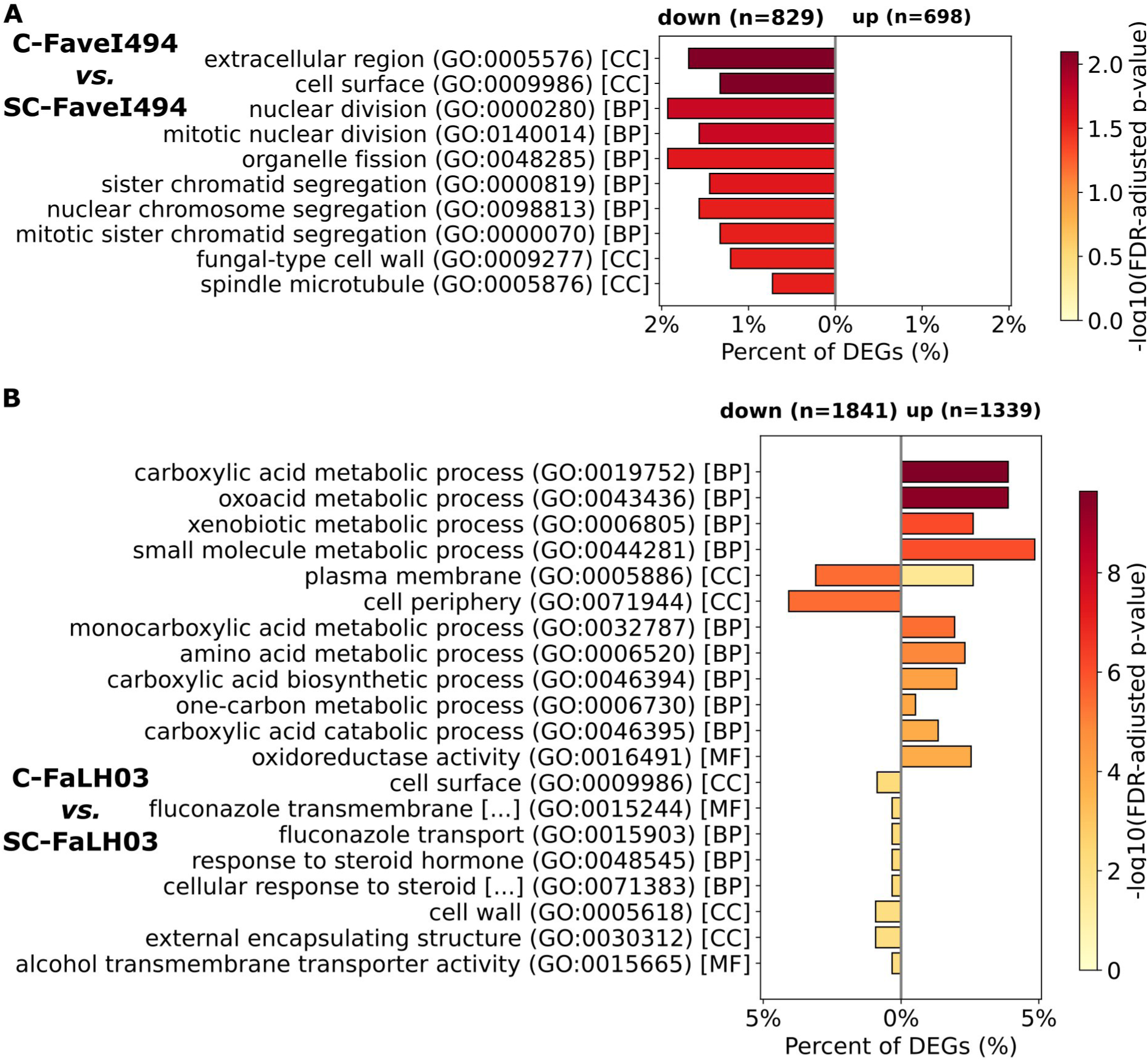
Gene Ontology (GO) term enrichment analysis of *F. avenaceum* differentially expressed genes in confrontation with FgrI349, relative to self-confrontation. **(A)** C-FaveI494 *vs*. SC-FaveI494. **(B)** C-FaLH03 *vs*. SC-FaLH03. Horizontal bar plots show enriched GO categories for up-regulated (right side) and down-regulated (left side) genes. The length of each bar indicates the percentage of differentially expressed genes annotated to the respective GO term, while color intensity represents the statistical significance of enrichment expressed as -Log₁₀(adjusted p-value).

## References

Anderson DW, Black RM, Lee CG, Pottage C, Rickard RL, Sandford MS, et al. Structure-activity studies of trichothecenes: cytotoxicity of analogs and reaction products derived from T-2 toxin and neosolaniol. J Med Chem. 1989 Mar 1;32(3):555–62. doi:10.1021/jm00123a008

Avalos J, Estrada AF. Regulation by light in Fusarium. Fungal Genet Biol. 2010 Nov;47(11):930–8. doi:10.1016/j.fgb.2010.05.001 PubMed PMID: 20460165.

Avalos J, Limón MC. Fungal Secondary Metabolism. Encyclopedia. 2022 Mar;2(1):1–13. doi:10.3390/encyclopedia2010001

Beccari G, Prodi A, Tini F, Bonciarelli U, Onofri A, Oueslati S, et al. Changes in the Fusarium Head Blight Complex of Malting Barley in a Three-Year Field Experiment in Italy. Toxins (Basel). 2017 Mar 29;9(4):120. doi:10.3390/toxins9040120 PubMed PMID: 28353653; PubMed Central PMCID: PMC5408194.

Bennett JW, Klich M. Mycotoxins. Clin Microbiol Rev. 2003 Jul;16(3):497–516. doi:10.1128/CMR.16.3.497-516.2003 PubMed PMID: 12857779; PubMed Central PMCID: PMC164220.

Brakhage AA. Regulation of fungal secondary metabolism. Nat Rev Microbiol. 2013 Jan;11(1):21–32. doi:10.1038/nrmicro2916 PubMed PMID: 23178386.

Buivydaitė Ž, Winding A, Jørgensen LN, Zervas A, Sapkota R. New insights into RNA mycoviruses of fungal pathogens causing Fusarium head blight. Virus Res. 2024 Sep 13;349:199462. doi:10.1016/j.virusres.2024.199462 PubMed PMID: 39260572; PubMed Central PMCID: PMC11417338.

Cho WK, Yu J, Lee KM, Son M, Min K, Lee YW, et al. Genome-wide expression profiling shows transcriptional reprogramming in Fusarium graminearum by Fusarium graminearum virus 1-DK21 infection. BMC Genomics. 2012a;13(1):173. doi:10.1186/1471-2164-13-173

Cho WK, Yu J, Lee KM, Son M, Min K, Lee YW, et al. Genome-wide expression profiling shows transcriptional reprogramming in Fusarium graminearum by Fusarium graminearum virus 1-DK21 infection. BMC Genomics. 2012b May 6;13:173. doi:10.1186/1471-2164-13-173 PubMed PMID: 22559730; PubMed Central PMCID: PMC3478160.

De Clerck C, Josselin L, Vangoethem V, Lassois L, Fauconnier ML, Jijakli H. Weapons against Themselves: Identification and Use of Quorum Sensing Volatile Molecules to Control Plant Pathogenic Fungi Growth. Microorganisms. 2022 Dec;10(12):2459. doi:10.3390/microorganisms10122459

Dweba CC, Figlan S, Shimelis HA, Motaung TE, Sydenham S, Mwadzingeni L, et al. Fusarium head blight of wheat: Pathogenesis and control strategies. Crop Protection. 2017 Jan 1;91:114–22. doi:10.1016/j.cropro.2016.10.002

Eagan JL, Keller NP. Fungal secondary metabolism. Current Biology. 2025 Jun 9;35(11):R503–8. doi:10.1016/j.cub.2025.02.029

Ederli L, Beccari G, Tini F, Bergamini I, Bellezza I, Romani R, et al. Enniatin b and deoxynivalenol activity on bread wheat and on fusarium species development [Internet]. 2021a. doi:10.3390/toxins13100728

Ederli L, Beccari G, Tini F, Bergamini I, Bellezza I, Romani R, et al. Enniatin B and Deoxynivalenol Activity on Bread Wheat and on Fusarium Species Development. Toxins. 2021b Oct;13(10):728. doi:10.3390/toxins13100728

Gentleman RC, Carey VJ, Bates DM, Bolstad B, Dettling M, Dudoit S, et al. Bioconductor: open software development for computational biology and bioinformatics. Genome Biol. 2004;5(10):R80. doi:10.1186/gb-2004-5-10-r80 PubMed PMID: 15461798; PubMed Central PMCID: PMC545600.

Glenn AE. Mycotoxigenic *Fusarium* species in animal feed. Animal Feed Science and Technology. 2007 Oct 1;Fusarium and their toxins: Mycology, occurrence, toxicity, control and economic impact137(3):213–40. doi:10.1016/j.anifeedsci.2007.06.003

Gluck-Thaler E, Ralston T, Konkel Z, Ocampos CG, Ganeshan VD, Dorrance AE, et al. Giant Starship Elements Mobilize Accessory Genes in Fungal Genomes. Mol Biol Evol. 2022 May 3;39(5):msac109. doi:10.1093/molbev/msac109 PubMed PMID: 35588244; PubMed Central PMCID: PMC9156397.

Huber W, Carey VJ, Gentleman R, Anders S, Carlson M, Carvalho BS, et al. Orchestrating high-throughput genomic analysis with Bioconductor. Nat Methods. 2015 Feb;12(2):115–21. doi:10.1038/nmeth.3252 PubMed PMID: 25633503; PubMed Central PMCID: PMC4509590.

Ichikawa T, Tanaka M, Watanabe T, Zhan S, Watanabe A, Shintani T, et al. Crucial role of the intracellular α-glucosidase MalT in the activation of the transcription factor AmyR essential for amylolytic gene expression in Aspergillus oryzae. Biosci Biotechnol Biochem. 2021 Sep 1;85(9):2076–83. doi:10.1093/bbb/zbab125

Inoue N, Tsuge K, Yanagita T, Oikawa A, Nagao K. Time-Course Metabolomic Analysis: Production of Betaine Structural Analogs by Fungal Fermentation of Seaweed. Metabolites. 2024 Apr 3;14(4):201. doi:10.3390/metabo14040201 PubMed PMID: 38668329; PubMed Central PMCID: PMC11051755.

Janik E, Niemcewicz M, Ceremuga M, Stela M, Saluk-Bijak J, Siadkowski A, et al. Molecular Aspects of Mycotoxins-A Serious Problem for Human Health. Int J Mol Sci. 2020 Oct 31;21(21):8187. doi:10.3390/ijms21218187 PubMed PMID: 33142955; PubMed Central PMCID: PMC7662353.

Jestoi M. Emerging fusarium-mycotoxins fusaproliferin, beauvericin, enniatins, and moniliformin: a review. Crit Rev Food Sci Nutr. 2008 Jan;48(1):21–49. doi:10.1080/10408390601062021 PubMed PMID: 18274964.

Keller NP. Fungal secondary metabolism: regulation, function and drug discovery. Nat Rev Microbiol. 2019;17(3):167–80. doi:10.1038/s41579-018-0121-1

Kelly AC, Ward TJ. Population genomics of Fusarium graminearum reveals signatures of divergent evolution within a major cereal pathogen. PLoS One. 2018;13(3):e0194616. doi:10.1371/journal.pone.0194616 PubMed PMID: 29584736; PubMed Central PMCID: PMC5870968.

King R, Urban M, Hammond-Kosack KE. Annotation of Fusarium graminearum (PH-1) Version 5.0. Genome Announc. 2017 Jan 12;5(2):e01479–16. doi:10.1128/genomeA.01479-16 PubMed PMID: 28082505; PubMed Central PMCID: PMC5256205.

King R, Urban M, Hammond-Kosack MCU, Hassani-Pak K, Hammond-Kosack KE. The completed genome sequence of the pathogenic ascomycete fungus Fusarium graminearum. BMC Genomics. 2015 Jul 22;16(1):544. doi:10.1186/s12864-015-1756-1 PubMed PMID: 26198851; PubMed Central PMCID: PMC4511438.

Kulik T, Jestoi M, Okorski A. Development of TaqMan assays for the quantitative detection of Fusarium avenaceum/Fusarium tricinctum and Fusarium poae esyn1 genotypes from cereal grain. FEMS Microbiol Lett. 2011 Jan;314(1):49–56. doi:10.1111/j.1574-6968.2010.02145.x PubMed PMID: 21059180.

Kuroki M, Yaguchi T, Urayama S ichi, Hagiwara D. Experimental verification of strain-dependent relationship between mycovirus and its fungal host. iScience. 2023;26(8):107337. doi:10.1016/j.isci.2023.107337

Lee KM, Cho WK, Yu J, Son M, Choi H, Min K, Lee YW, Kim KH. A comparison of transcriptional patterns and mycological phenotypes following infection of Fusarium graminearum by four mycoviruses. PLoS One. 2014;9(6):e100989. doi:10.1371/journal.pone.0100989 PubMed PMID: 24964178; PubMed Central PMCID: PMC4071046.

Lee KM, Cho WK, Yu J, Son M, Choi H, Min K, Lee YW, Kim KH, et al. A Comparison of Transcriptional Patterns and Mycological Phenotypes following Infection of Fusarium graminearum by Four Mycoviruses. Yu JH, editor. PLoS ONE. 2014 Jun 25;9(6):e100989. doi:10.1371/journal.pone.0100989

Liu GY, Nizet V. Color me bad: microbial pigments as virulence factors. Trends Microbiol. 2009 Sep;17(9):406–13. doi:10.1016/j.tim.2009.06.006 PubMed PMID: 19726196; PubMed Central PMCID: PMC2743764.

Lysøe E, Harris LJ, Walkowiak S, Subramaniam R, Divon HH, Riiser ES, et al. The genome of the generalist plant pathogen Fusarium avenaceum is enriched with genes involved in redox, signaling and secondary metabolism. PLoS One. 2014;9(11):e112703. doi:10.1371/journal.pone.0112703 PubMed PMID: 25409087; PubMed Central PMCID: PMC4237347.

Ma LJ, van der Does HC, Borkovich KA, Coleman JJ, Daboussi MJ, Di Pietro A, et al. Comparative genomics reveals mobile pathogenicity chromosomes in Fusarium. Nature. 2010 Mar;464(7287):367–73. doi:10.1038/nature08850

McGinnis MR. Laboratory handbook of medical mycology. New York: Academic Press; 1980. xiii, 661 p.

Nielsen LK, Jensen JD, Nielsen GC, Jensen JE, Spliid NH, Thomsen IK, et al. Fusarium Head Blight of Cereals in Denmark: Species Complex and Related Mycotoxins. Phytopathology®. 2011;101(8):960–9. doi:10.1094/PHYTO-07-10-0188

Osborne LE, Stein JM. Epidemiology of Fusarium head blight on small-grain cereals. Int J Food Microbiol. 2007 Oct 20;119(1–2):103–8. doi:10.1016/j.ijfoodmicro.2007.07.032 PubMed PMID: 17716761.

Pa E, A P, S F, H P, J A, A W, et al. The nf-core framework for community-curated bioinformatics pipelines. Nature biotechnology. 2020 Mar;38(3). doi:10.1038/s41587-020-0439-x PubMed PMID: 32055031.

Pang Z, Lu Y, Zhou G, Hui F, Xu L, Viau C, et al. MetaboAnalyst 6.0: towards a unified platform for metabolomics data processing, analysis and interpretation. Nucleic Acids Res. 2024 Jul 5;52(W1):W398–406. doi:10.1093/nar/gkae253

Parker D, Meyling NV, De Fine Licht HH. Phenotypic variation and genomic variation in insect virulence traits reveal patterns of intraspecific diversity in a locust-specific fungal pathogen. j evol Biol. 2023 Oct 1;36(10):1438–54. doi:10.1111/jeb.14214

Parry DW, Jenkinson P, McLEOD L. Fusarium ear blight (scab) in small grain cereals—a review. Plant Pathology. 1995;44(2):207–38. doi:10.1111/j.1365-3059.1995.tb02773.x

Patel H, Ewels P, Manning J, Garcia MU, Peltzer A, Hammarén R, et al. nf-core/rnaseq: nf-core/rnaseq v3.18.0 -Lithium Lynx [Internet]. Zenodo; 2024. Available from: https://doi.org/10.5281/zenodo.14537300 doi:10.5281/zenodo.14537300

Patro R, Duggal G, Love MI, Irizarry RA, Kingsford C. Salmon provides fast and bias-aware quantification of transcript expression. Nat Methods. 2017 Apr;14(4):417–9. doi:10.1038/nmeth.4197

Petrucci A, Khairullina A, Sarrocco S, Jensen DF, Jensen B, Jørgensen HJL, et al. Understanding the mechanisms underlying biological control of Fusarium diseases in cereals. Eur J Plant Pathol. 2023a Dec 1;167(4):453–76. doi:10.1007/s10658-023-02753-5

Petrucci A, Khairullina A, Sarrocco S, Jensen DF, Jensen B, Jørgensen HJL, et al. Understanding the mechanisms underlying biological control of Fusarium diseases in cereals. Eur J Plant Pathol. 2023b Dec 1;167(4):453–76. doi:10.1007/s10658-023-02753-5

Petrucci A, Vicente I, Cesarini M, Susca A, Sarrocco S, Vannacci G. Fusarium graminearum regulates kp4l genes, encoding killer toxins, during competitive interaction with other plant pathogenic Fusarium species. Fungal Biology. 2025 Jun 1;129(4):101569. doi:10.1016/j.funbio.2025.101569

Pfannenstiel BT, Keller NP. On top of biosynthetic gene clusters: How epigenetic machinery influences secondary metabolism in fungi. Biotechnol Adv. 2019 Nov 1;37(6):107345. doi:10.1016/j.biotechadv.2019.02.001 PubMed PMID: 30738111; PubMed Central PMCID: PMC6685777.

Ritchie ME, Phipson B, Wu D, Hu Y, Law CW, Shi W, et al. limma powers differential expression analyses for RNA-sequencing and microarray studies. Nucleic Acids Res. 2015 Apr 20;43(7):e47. doi:10.1093/nar/gkv007

Saravanakumar K, Yu C, Dou K, Wang M, Li Y, Chen J. Synergistic effect of *Trichoderma*-derived antifungal metabolites and cell wall degrading enzymes on enhanced biocontrol of *Fusarium oxysporum* f. sp. *cucumerinum*. Biological Control. 2016 Mar 1;94:37–46. doi:10.1016/j.biocontrol.2015.12.001

Schindelin J, Arganda-Carreras I, Frise E, Kaynig V, Longair M, Pietzsch T, et al. Fiji -an Open Source platform for biological image analysis. Nature methods. 2012 Jun 28;9(7):10.1038/nmeth.2019. doi:10.1038/nmeth.2019 PubMed PMID: 22743772.

Shore D, Albert B. Ribosome biogenesis and the cellular energy economy. Current Biology. 2022 Jun 20;32(12):R611–7. doi:10.1016/j.cub.2022.04.083

Smith JS, Thakur RA. Occurrence and fate of fumonisins in beef. Adv Exp Med Biol. 1996;392:39–55. doi:10.1007/978-1-4899-1379-1_4 PubMed PMID: 8850604.

Studt L, Wiemann P, Kleigrewe K, Humpf HU, Tudzynski B. Biosynthesis of fusarubins accounts for pigmentation of Fusarium fujikuroi perithecia. Appl Environ Microbiol. 2012 Jun;78(12):4468–80. doi:10.1128/AEM.00823-12 PubMed PMID: 22492438; PubMed Central PMCID: PMC3370568.

Takeshita N. Coordinated process of polarized growth in filamentous fungi. Biosci Biotechnol Biochem. 2016 Sep;80(9):1693–9. doi:10.1080/09168451.2016.1179092 PubMed PMID: 27121747.

Tsitsigiannis DI, Keller NP. Oxylipins as developmental and host-fungal communication signals. Trends Microbiol. 2007 Mar;15(3):109–18. doi:10.1016/j.tim.2007.01.005 PubMed PMID: 17276068.

Urbaniak M, Waśkiewicz A, Stępień Ł. Fusarium Cyclodepsipeptide Mycotoxins: Chemistry, Biosynthesis, and Occurrence. Toxins (Basel). 2020 Dec 3;12(12):765. doi:10.3390/toxins12120765 PubMed PMID: 33287253; PubMed Central PMCID: PMC7761704.

Wagacha JM, Oerke EC, Dehne HW, Steiner U. Interactions of Fusarium species during prepenetration development. Fungal Biology. 2012;116(7):836–47. doi:10.1016/j.funbio.2012.05.001

Wagacha JM, Steiner U, Dehne HW, Zuehlke S, Spiteller M, Muthomi J, et al. Diversity in Mycotoxins and Fungal Species Infecting Wheat in Nakuru District, Kenya. Journal of Phytopathology. 2010;158(7–8):527–35. doi:10.1111/j.1439-0434.2009.01653.x

Wang X, Bali M, Medintz I, Michels CA. Intracellular maltose is sufficient to induce MAL gene expression in Saccharomyces cerevisiae. Eukaryot Cell. 2002 Oct;1(5):696–703. doi:10.1128/EC.1.5.696-703.2002 PubMed PMID: 12455689; PubMed Central PMCID: PMC126750.

Windels CE. Economic and social impacts of fusarium head blight: changing farms and rural communities in the northern great plains. Phytopathology. 2000 Jan;90(1):17–21. doi:10.1094/PHYTO.2000.90.1.17 PubMed PMID: 18944567.

Xu XM, Nicholson P, Thomsett MA, Simpson D, Cooke BM, Doohan FM, et al. Relationship between the fungal complex causing Fusarium head blight of wheat and environmental conditions. Phytopathology. 2008 Jan 1;98(1):69–78. doi:10.1094/phyto-98-1-0069 PubMed PMID: 18943240.

Yu J, Kim KH. Exploration of the interactions between mycoviruses and Fusarium graminearum. In: Advances in Virus Research [Internet]. Academic Press; 2020 [cited 2025 Nov 8]. p. 123–44. Available from: https://www.sciencedirect.com/science/chapter/bookseries/abs/pii/S006535272030004Xdoi:10.1016/bs.aivir.2020.01.004

Zheng H, Kim J, Liew M, Yan JK, Herrera O, Bok JW, et al. Redox metabolites signal polymicrobial biofilm development via the napa oxidative stress cascade in aspergillus. Current Biology. 2015 Jan 5;25(1):29–37. doi:10.1016/j.cub.2014.11.018

